# Angiotensin II Type 1 Receptor Blockade Inhibits Gastric Cancer Metastasis Through Tight Junction Restoration

**DOI:** 10.64898/2026.01.08.698396

**Authors:** Sooraj Kakkat, Prabhat Suman, Sandeep Goswami, Brusi Kola, Kira A Bruno, Wendy L Frankel, Sujit Basu, Elba A Turbat-Herrera, Veronica Ramirez-Alcantara, Martin J Heslin, Joel F Andrews, Paramahansa Pramanik, Chandrani Sarkar, Debanjan Chakroborty

## Abstract

Cell-Cell adhesion maintained by tight junctions (TJs) is essential for epithelial integrity; loss of TJs correlates with poor prognosis, metastasis, and adverse clinical outcome in gastric cancer (GC). Restoring TJ integrity is therefore considered a promising therapeutic strategy in GC. The study identifies the stomach renin angiotensin system (stRAS) as a crucial regulator of TJ function in GC. Using integrative analysis of GC patient tissues, human GC cell lines, and orthotopic GC xenograft models, here we show that angiotensin II (ATII), the principal effector peptide of stRAS, drives TJ disassembly through an autocrine loop involving angiotensin receptor type 1 (AT1R) expressed on GC cells. Both ATII and AT1R are overexpressed in GC, where they suppress the expression of key TJ proteins. By analyzing global RNA-sequencing (RNA-seq) data and performing CRISPR/Cas9 gene deletion, chromatin immunoprecipitation, and functional assays, we mechanistically reveal that ATII, which is predominantly produced by cancer cells within the tumor microenvironment (TME), inhibits the expression of krüppel-like factor 4 (KLF4), a transcription factor crucial for the transcription of key TJ genes (*CLDN1, 3, 4*, and *TJP1*), leading to reduced synthesis of TJ proteins via AT1R expressed on cancer cells. Notably, the study demonstrates the effectiveness of pharmacological inhibition of AT1R with clinically established AT1R antagonists in preventing GC growth and metastasis by restoring TJ stability *in vivo*. These findings delineate a previously unrecognized role for ATII in governing TJ disassembly in GC and highlight the ATII/AT1R axis as a promising therapeutic target.

**GRAPHICAL ABSTRACT:** 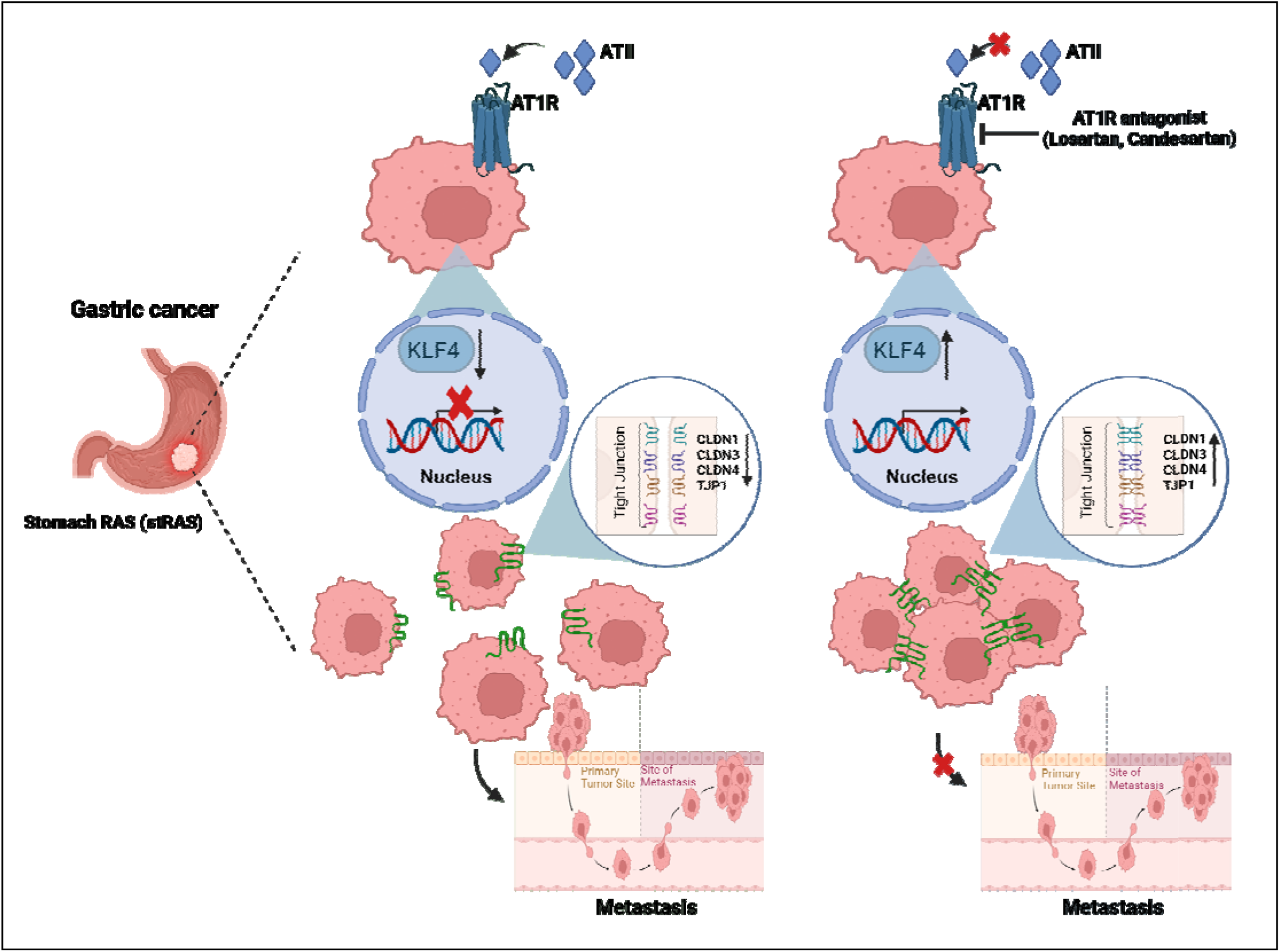

## INTRODUCTION

Stomach or gastric cancer (GC) is one of the most prevalent malignancies and a leading cause of cancer-related death in the world [1]. The disease is potentially curable when diagnosed early, with a survival rate of more than 90%. This number, however, sharply decreases to less than 20% when diagnosed at an advanced stage [2]. Newer advanced therapies have raised the bar for GC curability to ∼10-15% in resectable tumors. However, the overall 5-year survival rate for GC is still less than 25%, and the number has not changed in recent years because the majority of GC cases are diagnosed late, with more than 60% of patients presenting with metastatic disease [3]. Currently, the treatment options for these patients are limited, as surgery is less effective and patients often develop resistance to available agents quickly [2]. Therefore, there is a critical need for a better understanding of the process underlying GC metastasis to design novel, improved therapeutic strategies.

Metastasis initiation in solid tumors begins with cell-cell detachment within the primary tumor [4]. In their native environment, cells are held together by adhesive interactions and maintained by tight junctions (TJs), the disruption of which has been considered an early essential step in the cellular metastatic cascade [5]. TJs are primarily known for their role as tissue barriers, controlling the diffusion of selective molecules across paracellular spaces [6]. However, in epithelial cells, the primary function of TJs is to promote adhesive interactions and form tight attachments between adjacent cells [7]. Dysfunctional TJs, resulting from aberrant expression or distribution of TJ proteins, have been consistently linked to disease progression and a poorer prognosis in several epithelial cancers, including GC [8–12]. Several TJ components, including claudins, occludins, and zonula occludens, affect several aspects of tumorigenesis, including cellular differentiation, metastasis, and prognosis in GC. Nonetheless, the mechanisms that disrupt TJs in GC remain incompletely understood to date, creating a major roadblock to the design of effective strategies targeting TJ abnormalities in cancers. Therefore, gaining insight into the molecular mechanisms that regulate TJ disruption in GC is crucial, as this could lead to the development of potential therapeutic interventions to prevent the metastatic dissemination of cancer cells.

In this study, we reveal a novel role of locally expressed stomach tissue renin angiotensin system (stRAS) in initiating and facilitating GC cellular metastasis by disrupting TJ homeostasis between adjacent cancer cells within the primary tumor microenvironment. StRAS components, particularly Angiotensin II (ATII), the key effector peptide, and its receptor Angiotensin II receptor type 1 (AT1R), are significantly upregulated in GC compared to normal gastric tissues. Herein, we demonstrate that ATII, upon release from GC cells, interacts with AT1R expressed on GC cells to suppress krüppel-like factor 4 (KLF4), an evolutionarily conserved zinc finger transcription factor, and foster premetastatic niche development in GC. The suppression of KLF4 due to increased ATII-AT1R signaling in GC cells inhibits the synthesis of key TJ components, including claudins 1, 3, and 4 (CLDN1, 3, 4) and zonula occludens 1 (ZO-1), leading to dysfunctional TJs that facilitate cell detachment from their primary environment. Importantly, we identify that inhibition of AT1R is an effective strategy to prevent metastatic dissemination of GC cells, that reverses this cascade and restores junction integrity via KLF4 rescue in GC cells. The finding highlights the key role of the ATII-AT1R-KLF4 axis in mediating metastasis in GC and suggests that targeting this pathway may help prevent metastasis by restoring TJs in GC.

## Results

### Elevated expression of stomach renin angiotensin system, angiotensin II, and angiotensin II receptor type 1 in GC

To determine the role of stRAS in GC, we initially detected the expression of RAS components, ATII, and AT1R, in metastatic and non-metastatic human GC samples and normal stomach tissues collected from the same patients by immunohistochemistry (IHC) (Fig. 1A). Interestingly, a significant difference in the expression of ATII (Novus Biologicals, 1:100) was identified between normal, non-metastatic, and metastatic GC tissues collected from primary tumor sites, with metastatic tissues showing the highest expression of ATII within the TME (Fig. 1 A & B; *p*<0.05). AT1R expression was also significantly different between normal stomach and GC tissues (*p*<0.05); however, unlike ATII, the difference in the expression of AT1R (Abcam, 1:50) between non-metastatic and metastatic GC tissues did not reach a statistically significant value (Fig. 1 B; *p*=0.80). The above findings were subsequently validated in a larger cohort of GC patients by consulting the publicly available transcriptomic dataset from The University of Alabama at Birmingham Cancer Data Analysis portal (UALCAN) and The Cancer Genome Atlas (TCGA), which indicated a similar increase in the expression of *AGT* at the transcript level in human GC tissues compared to normal stomach tissues (Fig. 1C). The increased expressions of *AGT* and *AGTR1* also showed an association with reduced overall and disease-free survival in GC patients (Fig. 1D), suggesting the potential involvement of both of these stRAS components, in GC progression and outcome.

**Fig. 1.**
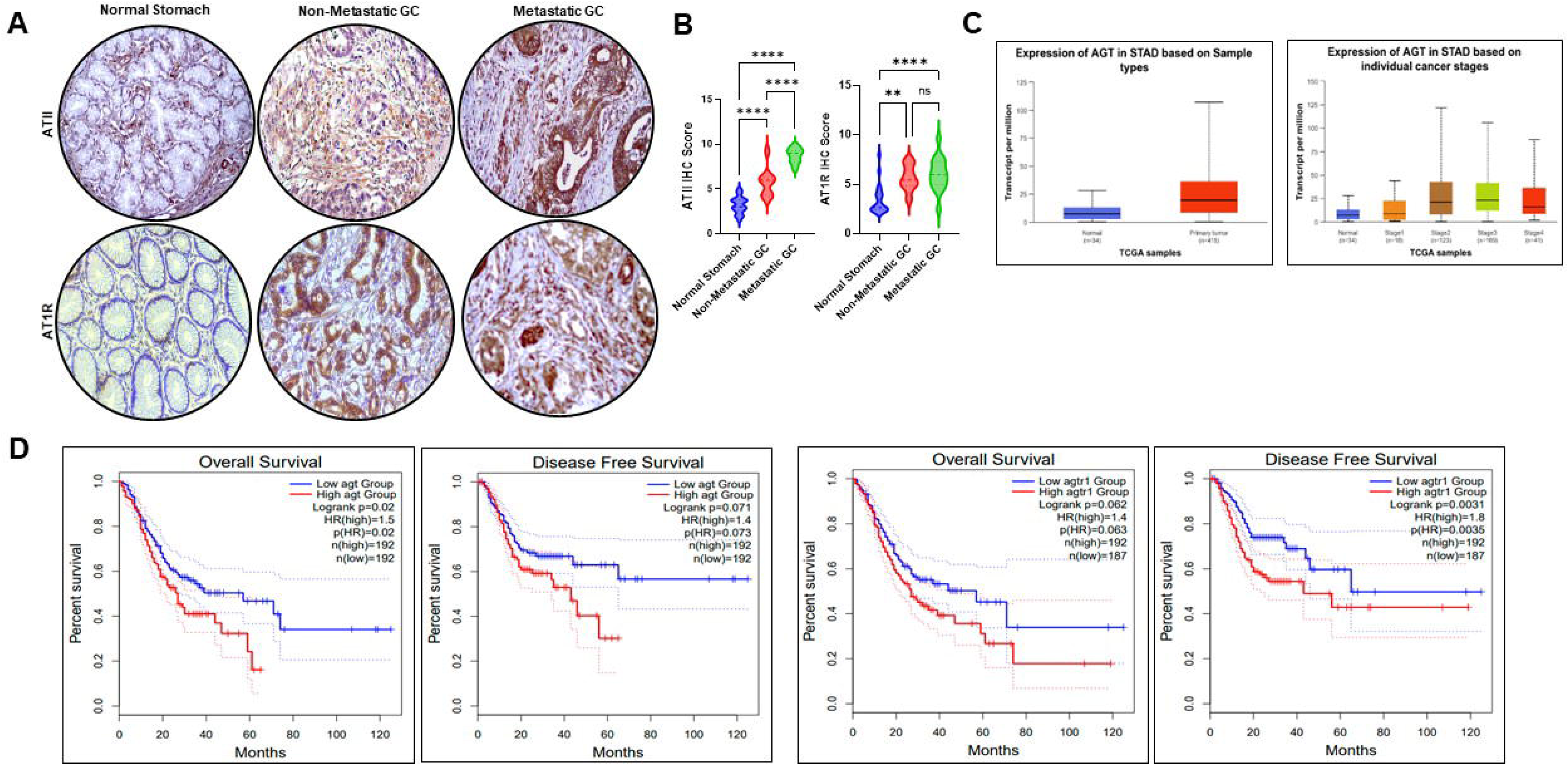
Increased ATII and AT1R expression in gastric cancer. **(A)** Representative IHC staining shows that ATII and its receptor, AT1R, are significantly elevated in GC tissues compared with normal stomach tissue. Magnification, 200X**. (B)** The violin plots show that the IHC scores for both ATII and AT1R are significantly higher in non-metastatic and metastatic GC tissues than in normal stomach tissue. Significance was determined by 1-way ANOVA (*n* = 64). (**C**) TCGA data show increased *AGT* expression in STAD compared to normal stomach, with a stage-wise increase in expression. (**D)** Kaplan-Meier survival curves from TCGA show OS and DFS for STAD patients with high and low *AGT* and *AGTR1* expression. High *AGT* expression is associated with significantly poor OS (*p =* 0.02), and high *AGTR1* expression is associated with significantly poor DFS (*p =* 0.0035). *\*\*P* < 0.01*; ****P* < 0.0001. ns: no significance. GC, gastric cancer; IHC, immunohistochemistry; STAD, stomach adenocarcinoma; OS, overall survival; DFS, disease-free survival; TCGA, the cancer genome atlas; AGT, angiotensin II; AGTR1, angiotensin II receptor type 1.

### Angiotensin II, acting through angiotensin II receptor type 1 expressed on GC cells, promotes GC cell metastasis

The next set of experiments was conducted to determine the source of ATII, the expression of AT1R, and the functional significance of ATII/AT1R overexpression in GC. For this, *in vitro* experiments were performed using human GC cell lines (MKN45, AGS, and NCI-N87). Results from the enzymatic immune assay (EIA, Sigma Aldrich), qRT-PCR, and western blot analysis identified GC cells as a potential source of ATII (Fig. 2A). Furthermore, western blot and qRT-PCR data indicated a strong expression of AT1R in these cancer cells (Fig. 2B). Therefore, to determine the functional role of ATII-AT1R in GC cells, ATII-secreting MKN45 and NCI-N87 GC cells were cultured to near confluence alone or in the presence of specific AT1R blockers /antagonists, losartan (1µM) and candesartan (1µM) [13,14]. Significant changes in GC cell growth and behavior were observed upon AT1R inhibition. Cell proliferation (Fig. 2C), along with migration and invasion (Fig. 2D and 2E), was significantly inhibited in losartan and candesartan-treated MKN45 and NCI-N87 cells compared to untreated GC cells. Taken together, the above results identified a novel autocrine function of the ATII/AT1R axis in mediating an aggressive behavior in GC cells.

**Fig. 2.**
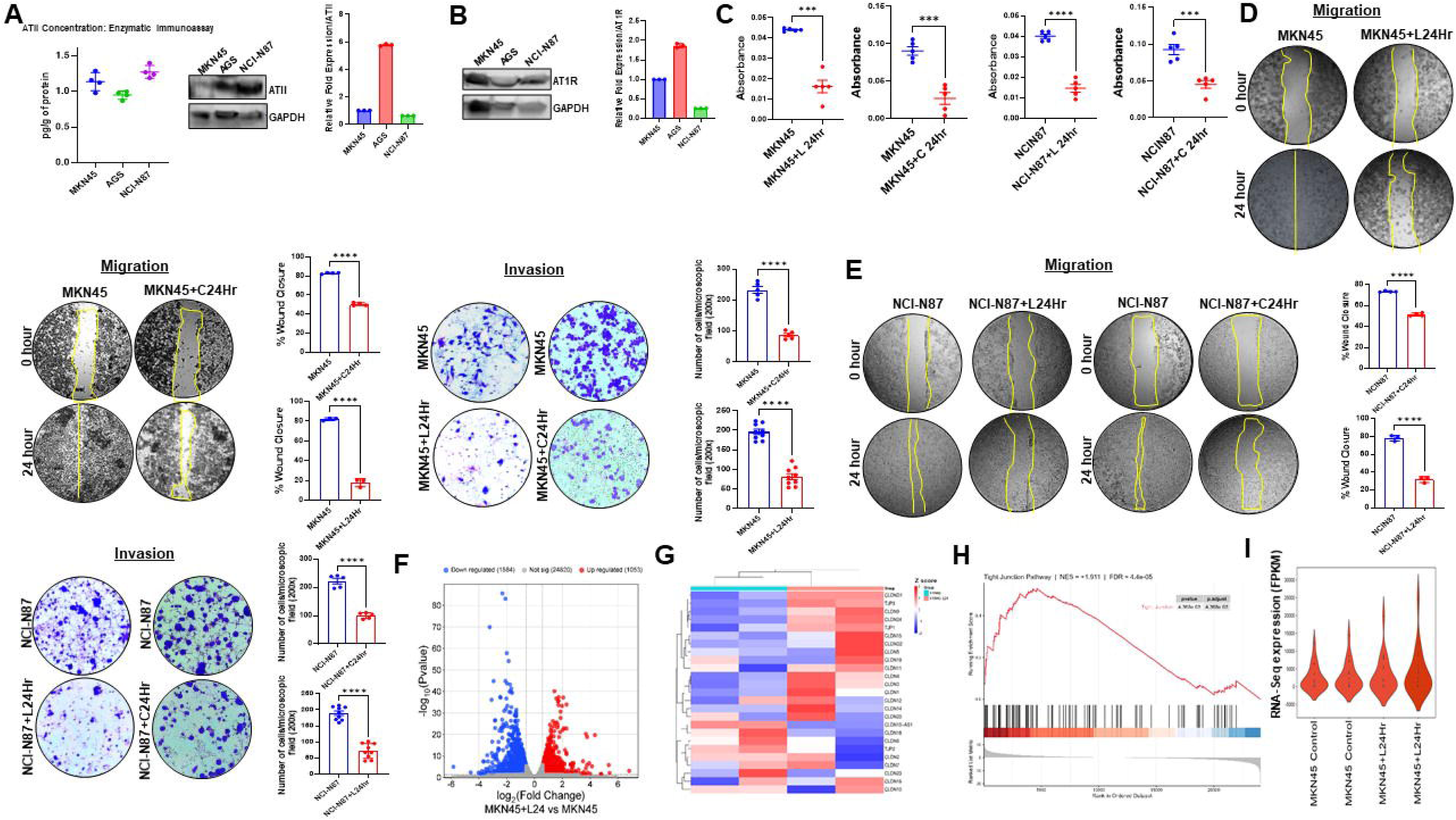
ATII promotes gastric cancer cell metastasis by regulating the expression of TJ proteins in gastric cancer cells via KLF4. **(A)** Enzymatic Immuno Assay (EIA), western blot, and qRT-PCR reveal expression of ATII in GC cells, MKN45, AGS, and NCI-N87. For quantification, the mRNA level of the ATII gene was normalized to the housekeeping gene GAPDH. Data represented as mean±SD of three independent experiments. **(B)** Western blot and qRT-PCR data show the expression of AT1R in GC cells. For quantification, the mRNA level of the ATIR gene was normalized to the housekeeping gene GAPDH. Data represented as mean±SD of three independent experiments. Blockade of AT1R by the AT1R antagonist losartan (1µM) and candesartan (1µM) significantly reduces GC cell **(C)** proliferation, **(D and E)** migration (magnification, 40X), and invasion (magnification, 200X). For the proliferation the data represented as mean±SEM and analyzed using 2-tailed *t* test (*n* = 5). The percentage of wound closure is shown in bar diagrams. Data represented as mean±SD and analyzed using 2-tailed *t* test (*n* = 3 for losartan treatment and *n* = 4 for candesartan treatment). Trans well assay shows that both losartan and candesartan treatment significantly reduces the invasive ability of MKN45 and NCI-N87 GC cells by inhibition of the AT1R receptors expressed in these cells. The number of invaded cells through the membrane in the control and AT1R antagonists treated groups is represented as a bar diagram. Data represented as mean±SEM and analyzed using 2-tailed *t* test (*n* = 9 for losartan treatment and *n* = 5 for candesartan treatment). (**F)** A volcano plot illustrating the differentially expressed genes (DEGs) obtained from RNA-sequencing (RNA-seq) analysis of losartan treated MKN45 GC cancer cells compared to control cells. Losartan treated GC cells with upregulated and downregulated genes are shown in red and blue colors respectively. (**G)** The heatmap illustrates the differential expression of TJ-related genes in losartan treated MKN45 cells compared to control cells. (**H)** GSEA plot demonstrates a positive enrichment of the TJ gene set in losartan treated MKN45 cells, suggesting that TJs are more intact upon treatment with losartan. **I**, Violin plot depicts the overall expression of TJ-related genes in losartan treated MKN45 cells compared with control cells. \*\*\**P* < 0.001, \*\*\*\**P* < 0.0001. GC, gastric cancer; EIA, enzymatic immunoassay; GSEA, gene set enrichment analysis; KEGG, kyoto encyclopedia of genes and genomes; TJ, tight junction.

### Inhibition of Angiotensin II receptor type 1 alters the tight junction gene signature in GC cells

Since significant changes in GC cell growth and metastasis were observed upon AT1R inhibition, subsequent experiments were conducted to identify the molecular events underlying these behavioral changes. For this, global RNA-sequencing (RNA-seq) was first performed. GC cells (MKN45) upon confluence were serum-starved overnight. The next day, cells were treated with the AT1R antagonist losartan (1 µM) or left untreated. Upon incubation, total RNA from both these groups of cells was extracted and deposited for sequencing and analysis (Novogene, Sacramento, CA, USA). RNA-seq was performed in duplicate on the Illumina NovaSeq 6000 using paired-end sequencing. Analysis of RNA-seq data derived from control and losartan treated MKN45 GC cells identified 2637 differentially expressed genes (DEGs) (fold change ≥1.5, p-value <0.05) between these two groups, of which 1053 were upregulated and 1584 downregulated in losartan treated cells compared to cells that were not treated (Fig. 2F). Among DEGs, a significant change in the expression of the genes associated with cell-cell attachments, cell polarity and TJ maintenance and formation were identified that showed significant upregulation in cells treated with losartan compared to untreated GC cells (Fig. 2G). The above observation was further validated by Gene Set Enrichment Analysis (GSEA), which revealed a significant positive enrichment of genes (NES=1.911, *p*-value=4.368e-05) (Fig. 2H) associated with TJ development and formation in the group of cells treated with losartan (Fig. 2I), suggesting the pivotal role of ATII in destabilizing TJ and facilitating the process of metastasis acting through AT1R expressed on these cells. The changes in TJ gene signature in GC cells upon AT1R blockade were further confirmed by qRT-PCR (Fig. 3A and B), followed by western blot analysis (Fig. 3C and D), and confocal microscopy (Fig. 3E and F), which clearly demonstrated a significant upregulation in the expression of key TJs, CLDN 1,3,4 and Zonula occludes 1 (ZO-1) both at transcript and protein level in MKN45 and NCI-N87 GC cells treated with losartan compared to cells that didn’t receive any treatment.

**Fig. 3.**
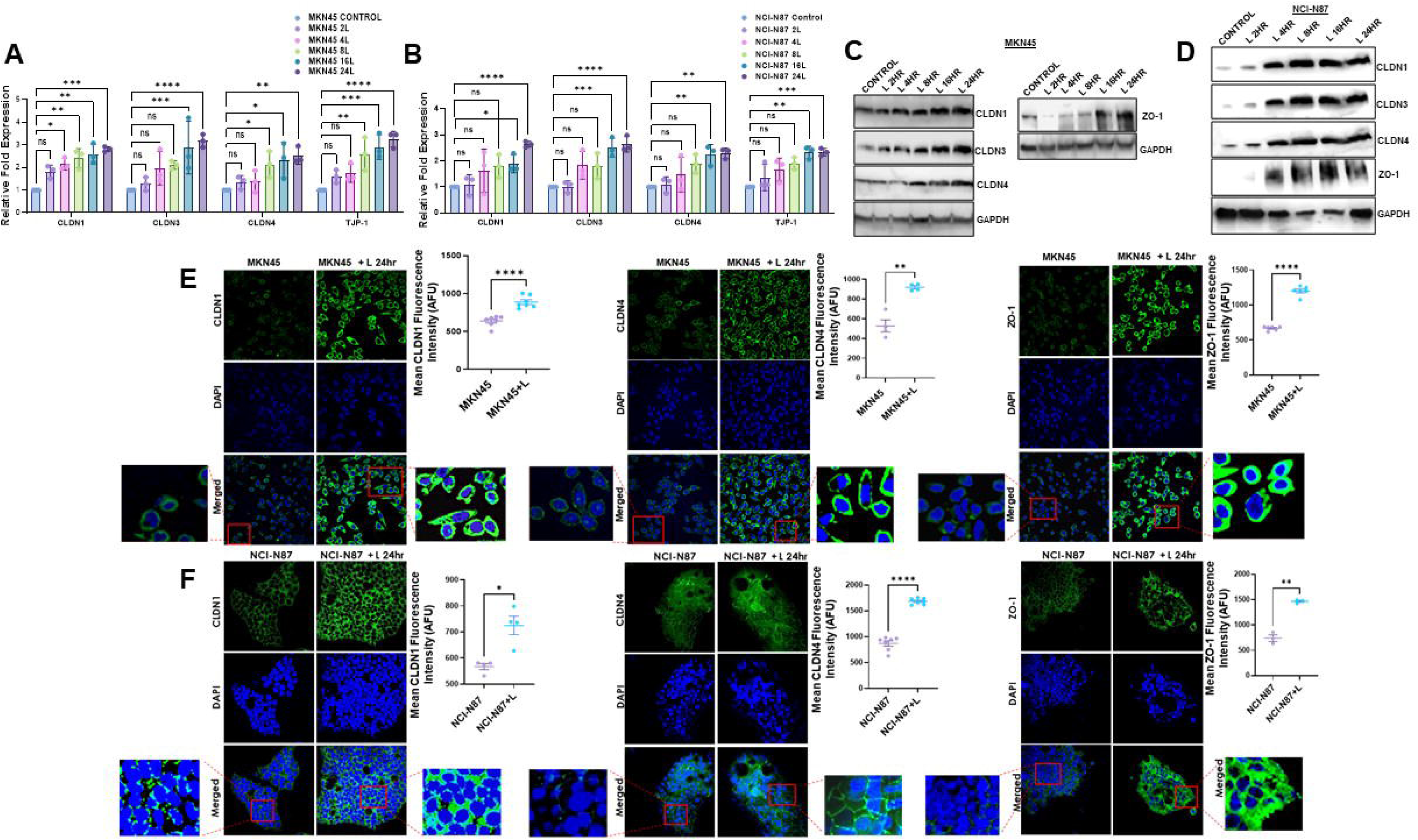
AT1R inhibition enhances the cell-cell cohesion in gastric cancer cells. (**A** and **B)** The qRT-PCR data show that blocking AT1R with losartan, over 2-24 hours, significantly increases the expression of TJ genes *CLDN1, CLDN3, CLDN4,* and *TJP1* in MKN45 and NCI-N87 GC cells. For quantification, mRNA levels of TJ genes were normalized to the housekeeping gene GAPDH. Data represented as mean±SD and analyzed using 2-way ANOVA with Dunnett’s multiple comparisons test (*n* = 3). (**C** and **D)** Western blot data also confirm increased expression of TJ proteins in GC cells upon AT1R blockade. (**E** and **F)** Immunofluorescence staining further demonstrates the enhanced cell-cell cohesion of Claudin 1, Claudin 4, and Zonula occludens 1 in losartan-treated GC cells compared to untreated controls. magnification, 200X. The green fluorescence intensity in maximum projection images of Z-stacks was quantified. The green channel was thresholded to create binary masks encompassing all cells, excluding background, and the mean intensity was measured in arbitrary fluorescence units (AFU), as shown in the bar diagram. In all images, the blue signal corresponds to DAPI-stained nuclei, and the green signal corresponds to tight junction proteins (Claudin 1, Claudin 4, and Zonula occludens 1) detected with Alexa Fluor-conjugated secondary antibodies. Data represented as mean±SEM of three independent experiments and analyzed using a 2-tailed t-test. \**P* < 0.05; \*\**P* <0.01;\*\*\**P* < 0.001; \*\*\*\**P* < 0.0001. *ns*: no significance. GC, gastric cancer; TJ, tight junction; AFU, arbitrary fluorescence units.

### Loss of tight junction proteins in human gastric cancer tissues strongly correlates with expression of angiotensin II and angiotensin II receptor type 1

The TJ status in human GC was next checked. For this, the expression of key TJ proteins was assessed in GC and nonmalignant adjacent tissues (normal stomach) collected from the same patients. Our IHC data indicated a significant loss of key TJ proteins, CLDN1, 3, 4 and ZO-1 in GC tissues compared to the normal stomach (Fig. 4A). Interestingly, a significant association was observed between ATII/AT1R expression and the expression of TJ proteins, CLDN 1, 3, 4, and ZO-1, where the expressions of ATII in human GC tissues was identified to be negatively correlated with the expressions of TJ proteins, i.e., CLDN 1 (*p*=0.0001), CLDN 3 (*p*<0.0001), CLDN 4 (*p*<0.0001) and ZO- 1 (*p*=0.0003) (Fig. 4B). Like ATII, the expression of AT1R was also negatively correlated with the expression of TJ proteins, i.e., CLDN 1 (*p*=0.0480), CLDN3 (*p*=0.0338), CLDN 4 (*p*=0.0267) and ZO-1 (*p*=0.3129) (Fig. 4C). The above association was further confirmed using RNA-seq data collected from TCGA-STAD cohort. For this, GC patients were stratified into low- and high- *AGT/AGTR1* expression groups using the cohort median as the cutoff (n = 100 per group for each gene). Data showed a clear gradient in the expression of TJ-related genes between high- and low- *AGT/AGTR1*-expressing GC groups. While GC patients with high *AGT/AGTR1* expression exhibited a disrupted signature (characterized by downregulation of key TJ- associated genes), a relatively improved TJ signature was noted in GC patients expressing low *AGT/AGTR1* (Fig. S1A). Taken together, these results indicate a strong connection between ATII/AT1R expression and TJ alterations in GC and suggest their possible role in advancing the disease.

**Fig. 4.**
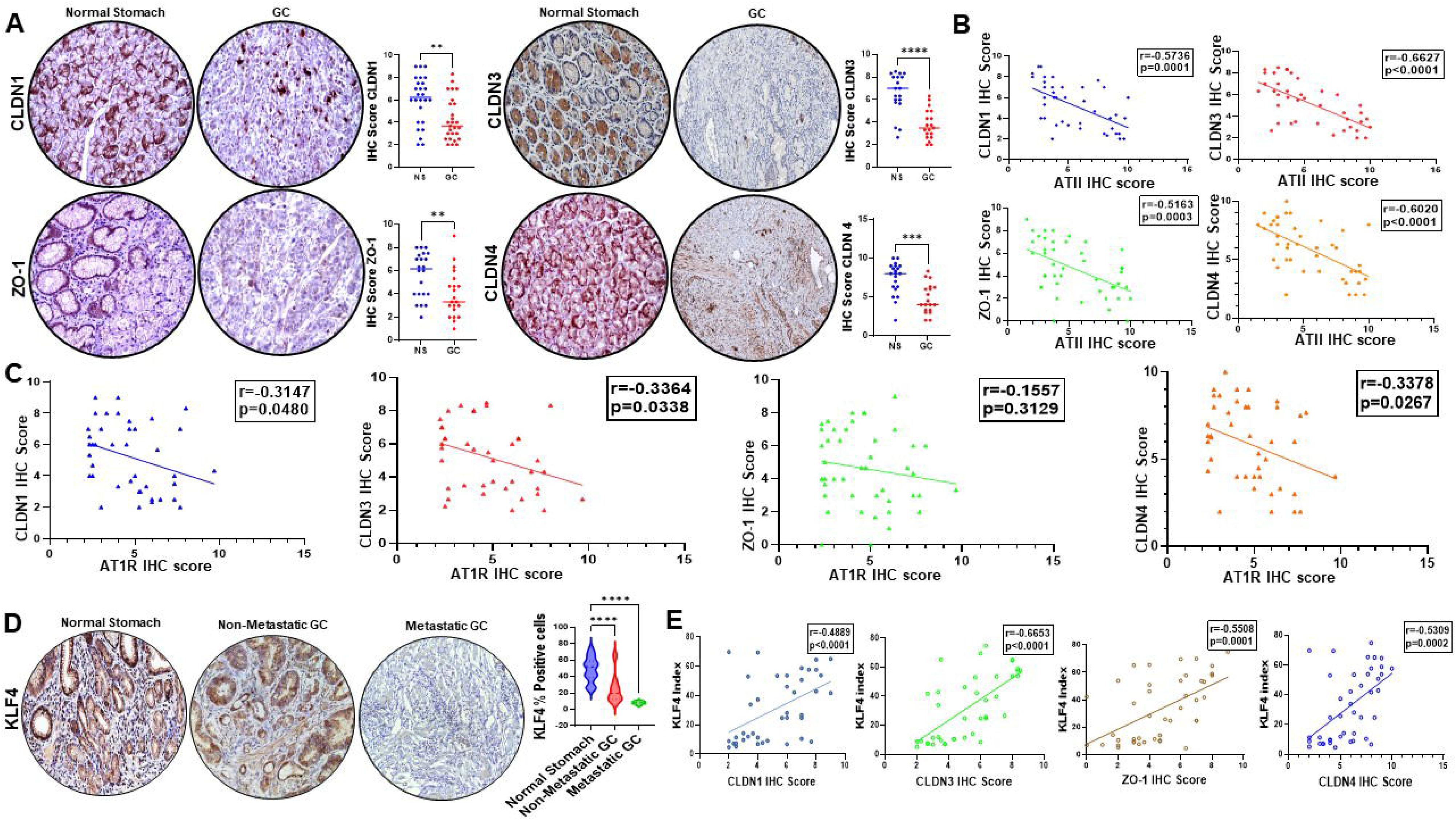
**Loss of tight junction proteins negatively correlates with ATII/AT1R expression and GC progression**. **(A)** IHC staining shows that the expression of TJ proteins Claudin 1, 3, 4, and Zonula occludens 1 is markedly decreased in human GC tissues compared to the normal stomach. The IHC score was analyzed and shown by the dot plot comparing IHC scores for Claudin 1 (n = 26 for NS; n = 27 for GC), 3 (n = 19 for NS; n = 19 for GC), 4 (n = 21 for NS; n = 19 for GC), and Zonula occludens 1 (n = 22 for NS; n = 21 for GC) in two groups: NS and GC. magnification, 200X. The IHC score for each TJ protein is significantly higher in normal stomach tissues compared to GC tissues. Each dot represents an individual data point, and the horizontal lines represent the median for each group. Data analyzed using a 2-tailed t-test. **(B)** Expression of ATII in human GC tissues shows an inverse correlation with the expression of TJ proteins, Claudin 1 (n = 40), 3 (n = 40), 4 (n = 43), and Zonula occludens 1 (n = 44). Correlation analysis between ATII expression and Claudins was performed using Pearson’s correlation test; the results are displayed in the graph. **(C)** Expression of AT1R in human GC tissues shows an inverse correlation with the expression of TJ proteins, Claudin 1 (n = 40), 3 (n = 40), 4 (n = 43), and Zonula occludens 1 (n = 44). Correlation analysis between ATIR expression and Claudins was performed using Pearson’s correlation test, and the values are displayed on the graph. **(D)** IHC images and quantification of KLF4-positive cells indicate a gradual loss of KLF4 expression in GC tissues with disease progression. Magnification, 200X. Significance was determined by 1-way ANOVA (n = 64). **(E)** Quantitative analysis reveals a strong positive correlation between the percentage of KLF4-positive cells and the expression of these TJ proteins, Claudin 1 (n = 40), 3 (n = 40), 4 (n = 43), and Zonula occludens 1 (n = 43) in GC tissues. Correlation analysis between KLF4 expression and Claudins was performed using Pearson’s correlation test, and the values are displayed on the graph. *\*\*P < 0.01;***P < 0.001; ****P < 0.0001*. ns: no significance. GC, gastric cancer; TJ, tight junction; NS, normal stomach; IHC, immunohistochemistry.

### Loss of Krüppel-like factor 4 in gastric cancer

Next, we sought to determine the underlying molecular mechanism by which ATII/AT1R mediates TJ disruption in GC. Reports, including those from our group, indicate that members of the KLF family, a zinc-finger transcription factor, play a critical role in regulating cell-cell integrity and barrier function across different cell and tissue types [15–17]. To determine the underlying molecular events leading to ATII-AT1Rlzlmediated TJ disruption in GC, we investigated the potential involvement of the KLF family of transcription factors, known for their involvement in biological processes like regulation of apoptosis, proliferation, cellular differentiation, preserving epithelial barrier integrity, and tissue architecture across different organs and tissues [18–21].

Interrogation and analysis of publicly available transcriptomic datasets from UALCAN identified alterations in the expression of several members of the KLF family transcription factor in human GC (Fig. S2A). However, a consistent trend was only noted in the expression of KLF4 and KLF5. While the expression of KLF5 was upregulated, the expression of KLF4 exhibited a particular pattern of downregulation that became more pronounced with the progression of GC (Fig. S2B). The above observation was confirmed further by consulting the protein expression database, Human Protein Atlas (Fig. S2C) [22], and immunohistochemical (IHC) analysis of GC tissues at different stages (non-metastatic and metastatic) collected from the department of pathology tissue archive at the Ohio State University and University of South Alabama Health Biobank.

Consistent with previous reports identifying KLF5 as a pro-proliferative transcription factor in most cancers, including GC, this observation reaffirms its pro-tumorigenic role in GC [18]. However, the role of KLF4 in cancers is conflicting, with both pro- and anti-tumorigenic effects that depend on the cancer type [19,23]. Our IHC data revealed a significant loss of KLF4-positive cells in GC tissues compared with normal stomach tissues isolated from the same patients, with the loss more pronounced in metastatic GC tissues (Fig. 4D). KLF4 plays diverse roles in cellular physiology. In addition to its prominent role in cell pluripotency, plasticity, and cell stemness, KLF4 in epithelial cells has been identified for its key role in keeping architectural integrity in the cell layer by maintaining cell-cell cohesion; loss of which compromises tissue integrity and has been considered a key initial step in the cellular metastatic cascade [20].

### Loss of Krüppel-like factor 4 expression in human gastric cancer tissues strongly correlates with the loss of tight junction proteins

The association between KLF4 loss and TJ status in GC tissues was checked next. For this, IHC scores of TJ proteins were correlated with the number of KLF4-positive cells. Our data identified a strong and positive correlation between the expressions of CLDN 1, 3, 4, and ZO- 1 proteins with the number of KLF4-positive cells in gastric tissues (Fig. 4E). These findings indicate the potential role of KLF4 in maintaining TJ integrity in gastric epithelial cells and suggest that its loss in GC might be a determining factor in initiating metastasis via TJ disintegration.

### Loss of Krüppel-like factor 4 facilitates gastric cancer cell metastasis

Analysis of available transcriptomic data and our IHC results revealed a significant loss of KLF4 during disease progression in GC. We next checked the expression status of KLF4 in different human GC cell lines (MKN45, AGS, and NCI-N87). Interestingly, our qRT-PCR and western blot data showed that all GC cells expressed KLF4 at varying levels, with MKN45 and NCI-N87 cells exhibiting significant KLF4 expression at both mRNA and protein levels (Fig. S3A). To simulate the KLF4 loss observed in GC patients, we next manipulated the expression of *KLF4* in GC cells (MKN45 and NCI-N87). The CRISPR/Cas9 gene deletion method was used to create *KLF4* knockout (KLF4KO) GC cell lines, KLF4KOMKN45 and KLF4KONCI-N87 (Fig. S3B and C). During generation, we also included a scramble control gRNA (Scr) as a negative control. Along with cell growth and proliferation (determined by doubling time) (Fig. S3D), our results showed a significant increase in the migration and invasion of GC cells upon *KLF4-*deletion compared to the control cells (Fig. S3E), identifying a potent tumor suppressive role of *KLF4* in gastric tumorigenesis.

### Loss of Krüppel-like factor 4 alters the expression of early and late metastatic genes in gastric cancer cells

Next, we investigated the molecular connection between KLF4 loss and GC cell metastasis. This was performed using MKN45 GC cells. Analyses of global RNA-Seq data from control MKN45 GC cells and KLF4KOMKN45 GC cells revealed a significant alteration in the expression of several early and late metastatic genes upon *KLF4*-deletion. RNA-seq of MKN45 GC cells and KLF4KOMKN45 GC cells identified 4188 DEGs (fold change ≥1.5, *p-*value <0.05), of which 1927 were upregulated and 2261 downregulated in KLF4KOMKN45 cells. The gene ontology and pathways of differentially expressed genes (DEGs) were analyzed with BigOmics Analytics [24]. The raw data were converted into counts per million and log2, and quantile normalization was applied. Other unwanted variations were removed using surrogate variable analysis in the sva package. The KEGG overrepresentation analysis of differentially expressed genes in *KLF4*-deleted cells identified a number of pathways that are enriched including the TJ pathway (*p*= 0.0009) (Fig. 5A). Additionally, gene ontology biological process analysis revealed that pathways related to miRNA mediated gene silencing were positively and significantly enriched in KLF4KOMKN45 cells, suggesting a potential role of *KLF4* in transcriptional regulation of TJ genes in GC cells (*p*= 0.0057) (Fig. 5B).

**Fig. 5.**
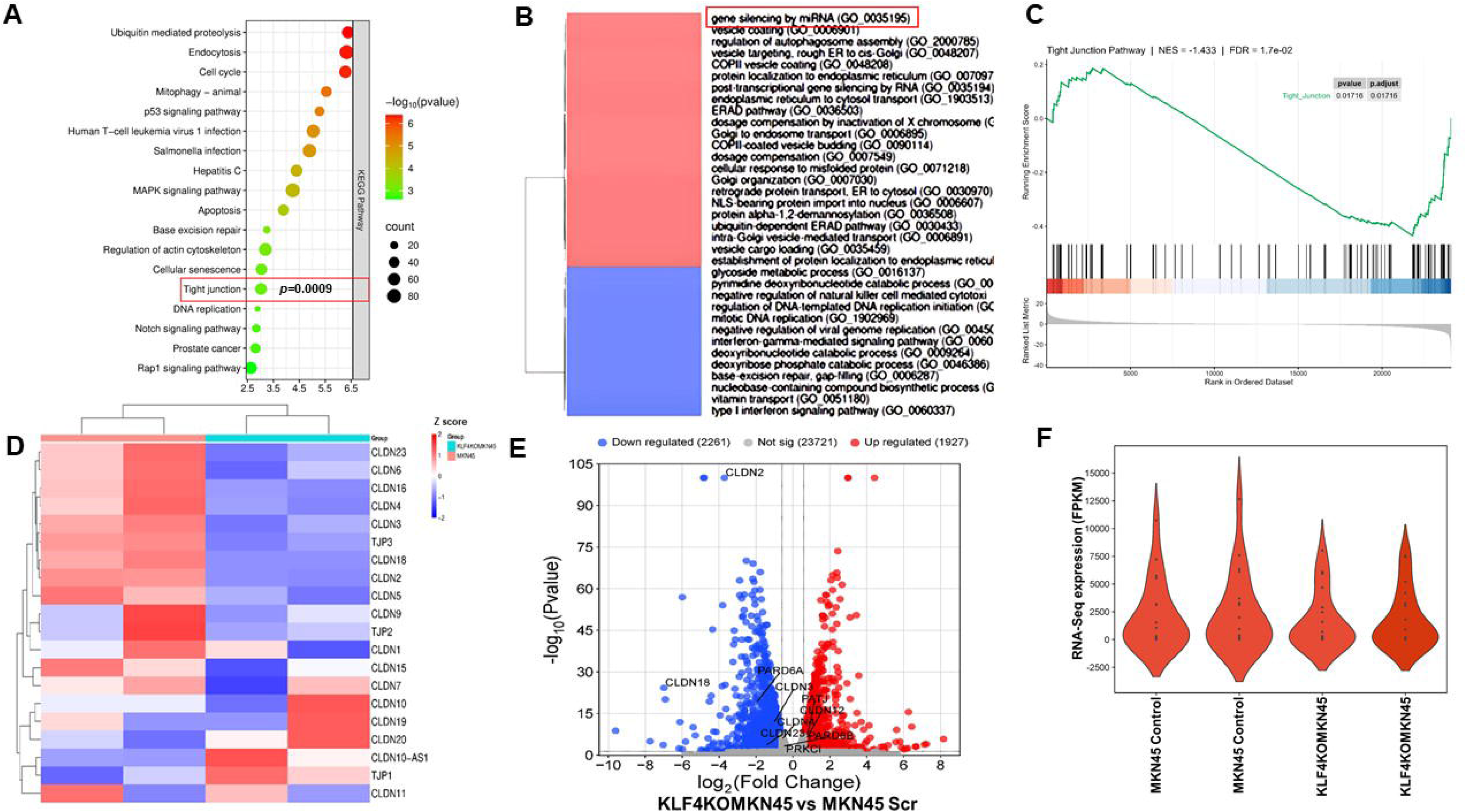
Loss of KLF4 results in the transcriptional repression of tight junction-related genes in GC cells. **(A)** KEGG overrepresentation analysis of differentially expressed genes in KLF4KO gastric cancer cells. Many pathways along with tight junction are significantly enriched in KLF4KO cells, indicating that these pathways are strongly affected by loss of KLF4. (**B)** Gene ontology biological process analysis of differentially expressed genes reveals a marked upregulation of pathways involved in miRNA-mediated gene silencing in KLF4KO cells. (**C)** Preranked GSEA of the full transcriptome using the tight junction gene set. The tight junction pathway shows a significant negative normalized enrichment score, indicating coordinated downregulation of tight junction-associated genes in KLF4KO cells compared with controls. (**D)** The heat map shows the differential expression of tight junction-related genes in KLF4KOMKN45 cells versus control cells; downregulation is particularly prominent across the claudin gene family. (**E)** Volcano plot highlights significantly downregulated tight junction genes (e.g., *CLDN2*, *CLDN3*, *CLDN4*, *PARD6A*, *CLDN23*, *CLDN18*, and *PRKCI*) and upregulated genes (e.g., *CLDN12*, *PATJ*) in KLF4KO GC cells. KLF4KOMKN45 GC cells with upregulated and downregulated genes are shown in red and blue colors, respectively. **(F)** The violin plot depicts the overall reduction in the expression of tight junction-related genes in KLF4KO cells compared with wild-type parental cells, consistent with impaired epithelial integrity following KLF4 loss. GC, gastric cancer; TJ, tight junction; GSEA, gene set enrichment analysis; KEGG, kyoto encyclopedia of genes and genomes.

Furthermore, Gene Set Enrichment Analysis revealed distinct divergence in TJ gene set enrichment in KLF4KOMKN45 cells compared to the control cells. The preranked GSEA analysis by using full ranked gene list showed a significant negative enrichment of TJ gene set, consistent with a corresponding downregulation of TJ components in *KLF4-*deleted cells (NES= -1.433, *p*-value=1.7e-02) (Fig. 5C). Differential expression analysis revealed that multiple genes involved in TJ formation and several members of the CLDN family, including *CLDN2*, *CLDN3*, *CLDN4*, *CLDN23*, and *CLDN18*, along with *PARD6A*, were significantly downregulated in KLF4KOMKN45 cells (Fig. 5D and E) compared to the control cells. Overall, the data identified significant downregulation of TJ-related genes, accompanied by increased cell metastasis in KLF4KO GC cells, indicating that KLF4 loss in GC is a key factor contributing to metastasis through TJ disruption between adjacent GC cells (Fig. 5F).

Western blot analyses (Fig. 6A), qRT-PCR (Fig. 6B), and confocal microscopy (Fig. 6C) further confirmed significant loss of important TJs, CLDN 1, 3, 4, and ZO-1 in both KLF4KOMKN45 and KLF4KONCI-N87 cells compared to scramble cells [25,26].

**Fig. 6.**
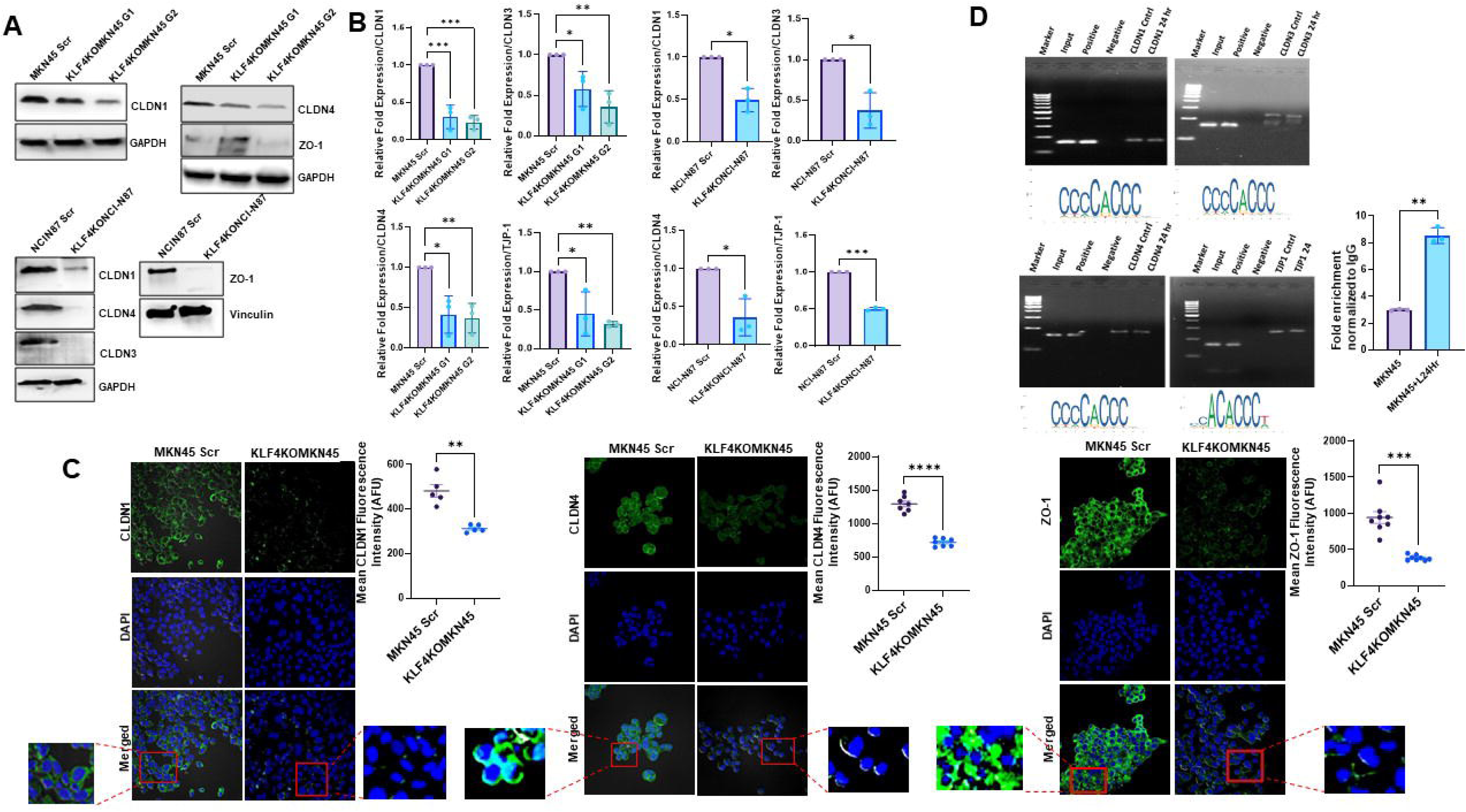
KLF4 regulates the expression of tight junction proteins in gastric cancer cells. **(A)** Western Blot analysis shows reduced expression of multiple TJ proteins, including Claudin 1, 3, 4, and Zonula occludens 1, in KLF4KOMKN45 and KLF4KONCI-N87 GC cells compared to scramble controls. **(B)** qRT-PCR analysis indicates significant downregulation of TJ genes, *CLDN1, CLDN3, CLDN4,* and *TJP1* at the transcript level in both KLF4KOMKN45 and KLF4KONCI-N87 cells. The quantification of mRNA levels of TJ genes was normalized to the housekeeping gene GAPDH. Data represented as mean±SD and analyzed using 1-way ANOVA with Dunnett’s multiple comparisons test and 2-tailed *t* test (n = 3) **(C)** Representative confocal images of immunofluorescence staining for TJ proteins in scrambled control and KLF4KOMKN45 cells. DAPI (Blue) is used as a nuclear stain, and the green signal corresponds to tight junction proteins (Claudin 1, Claudin 4, and Zonula occludens 1). Magnification, 200X. Loss of membrane-localized TJ proteins is evident in KLF4KO cells, indicating disruption of junctional integrity. The green fluorescence intensity in maximum projection images of Z-stacks was quantified. The green channel was thresholded to create binary masks encompassing all cells, excluding the background, and the mean intensity was measured in AFU, as shown in the bar diagram. Data represented as mean±SEM of three independent experiments and analyzed using a 2-tailed t-test. **(D)** ChIP followed by PCR demonstrates direct binding of KLF4 to the promoters of *CLDN1*, *CLDN3*, *CLDN4*, and *TJP1*, indicating transcriptional regulation of TJ genes by KLF4. *\*P* < 0.05; *\*\*P* < 0.01; *\*\*\*P* < 0.001; *\*\*\*\*P* < 0.0001. GC, gastric cancer; TJ, tight junction; ChIP, chromatin immunoprecipitation; Scr, scrambled control; AFU, arbitrary fluorescence units.

Since KEGG pathway analyses revealed changes in transcriptional control between wild-type and KLF4KO GC cells, and our data identified reduced TJ expression in KLF4KO GC cells, we next determined whether KLF4 transcriptionally regulates TJ gene expression. The JASPAR database was used to identify *KLF4* binding sites in the promoters of *CLDN 1*, *3*, *4*, and *TJP1*. The relatively higher binding score, indicating a stronger likelihood of *KLF4* binding, was then selected for further investigation. A ChIP assay was performed, and interestingly, our data revealed strong *KLF4* binding to promoters of *CLDN 1*, *3*, *4*, and *TJP1* (Fig. 6D).

### Elevated expressions of angiotensin II and angiotensin II receptor type 1 negatively regulate Krüppel-like factor 4 expression in gastric cancer

Next, we sought to identify the upstream events that prompt KLF4 loss or suppression in GC. In recent years, KLF4 has attracted significant attention as a crucial regulator of cardiac cellular hypertrophy. Interestingly, reports also indicate that ATII, by regulating KLF4 expression in cardiomyocytes, plays an important role in cardiac hypertrophy [27,28]. Given the high ATII and AT1R expression in GC tissues, we next investigated the role of the ATII/AT1R axis in regulating KLF4 expression. For this, the average IHC score of ATII/AT1R expression in individual tissues was correlated with the number of KLF4-positive cells in the same tissue. A significant association was observed between ATII/AT1R expression and the number of KLF4-positive cells, with ATII and AT1R negatively correlating with the number of KLF4-positive cells in GC tissues (Fig. S4A). This finding was also validated using transcriptomic data from cBioPortal, which showed a negative correlation between *AGT*/*AGTR1* and *KLF4* expression in GC (Fig. S4B)[29]. Next, the effect of ATII on KLF4 expression in GC cells was examined *in vitro*. Our results showed a significant increase in KLF4 expression in GC cells upon losartan treatment. Western blot data revealed a significant increase in KLF4 expression and nuclear enrichment over time in both MKN45 and NCI-N87 GC cells upon losartan treatment (Fig. 7A and B), which was further validated by confocal immunofluorescence analysis (Fig. 7C and D). Confocal microscopy images demonstrated a significant elevation in expression and nuclear incorporation, beginning at 2 hours after losartan treatment and gradually increasing up to 8 hours. Collectively, the results indicated a significant and prominent role of the ATII/AT1R in regulating KLF4 expression and function in GC cells.

**Fig. 7.**
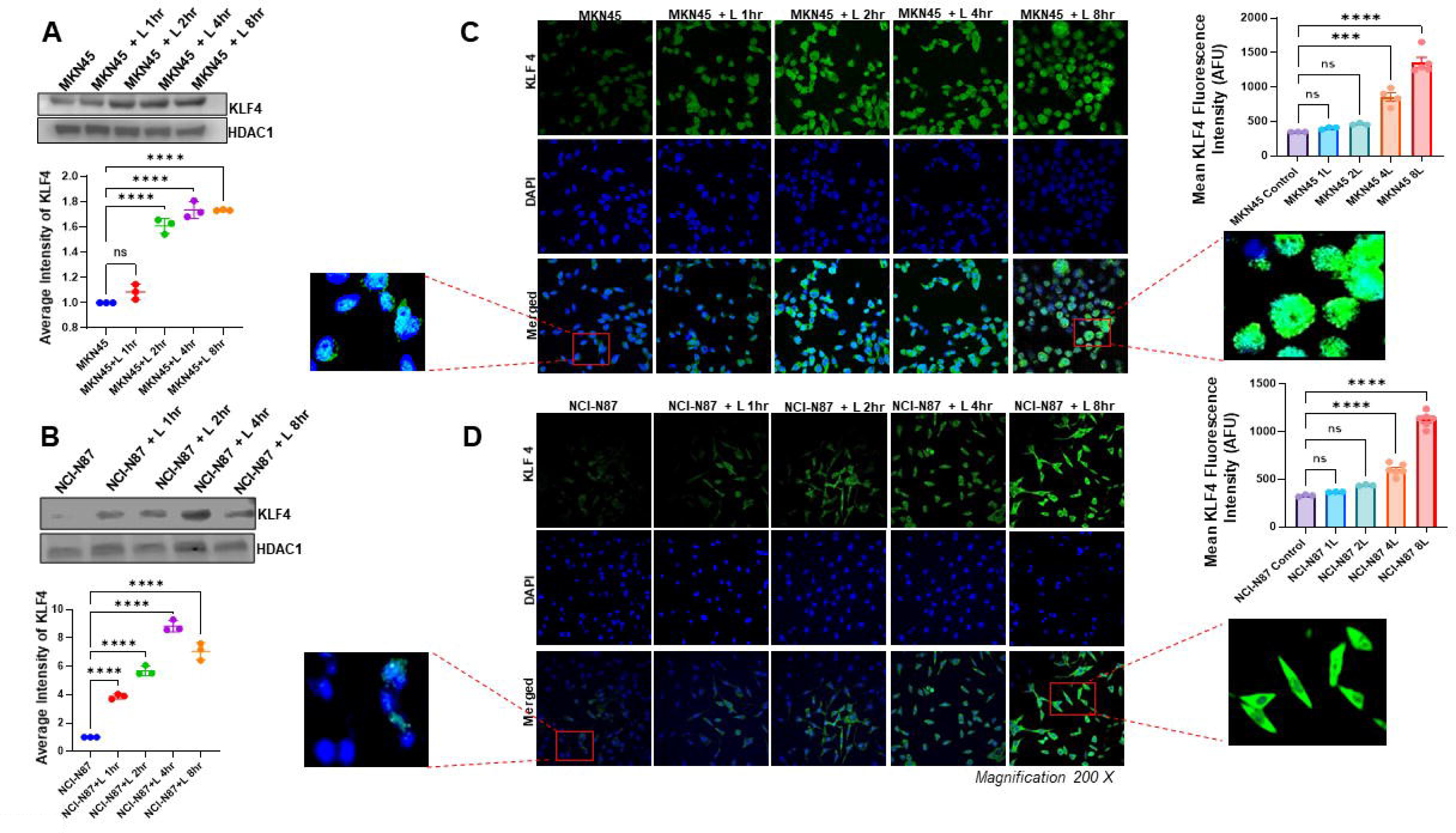
AT1R blockade induces KLF4 expression and nuclear localization in GC cells. (A and. **B)** Western blot data and densitometry analysis show a significant increase in KLF4 expression over time following losartan treatment of MKN45 and NCI-N87 GC cells. Data represented as mean±SD. Significance was determined by 1-way ANOVA with Dunnett’s multiple comparisons test. The densitometric quantification of KLF4 protein levels normalized to HDAC1 as a loading control. **(C and D)** Confocal microscopy also confirms nuclear enrichment of KLF4 in MKN45 and NCI-N87 cells upon losartan treatment. DAPI (Blue) is used as a nuclear stain, and the green signal corresponds to KLF4. Magnification, 200X. The green fluorescence intensity in maximum projection images of Z-stacks was quantified. The green channel was thresholded to create binary masks encompassing all cells, excluding the background, and the mean intensity was measured in arbitrary fluorescence units, as shown in the bar diagram. Data represented as mean±SEM. Significance was determined by 1-way ANOVA with Dunnett’s multiple comparisons test. *\*\*\*P <* 0.001*; ****P <* 0.0001. GC, gastric cancer.

### Angiotensin II receptor type 1 inhibition fails to restore tight junction integrity to suppress gastric cancer cell metastasis in the absence of Krüppel-like factor 4

Our next set of experiments was conducted to confirm that ATII/AT1R-mediated TJ disruption was through KLF4. For this, KLF4KOMKN45 and KLF4KONCI-N87 cells were used. Cell migration, invasion, and TJ protein expression were checked upon losartan treatment and compared with those of untreated cells. Results from scratch wound healing assays and cell invasion assays indicated that in the absence of KLF4 in GC cells, losartan treatment has no significant impact on inhibiting cell invasion and migration, as the losartan-treated cells, compared to control cells, exhibited comparable rates of wound closure over the observed period (24Hr) (Fig. S5A and B). Similarly, we observed that treatment with another AT1R blocker, candesartan, also had no significant effect on cell invasion or migration in the absence of KLF4 (Fig. S5C and D). Quantitative analysis also showed that the wound closure and invasion index were similar between the two groups of cells, with no significant difference in migration or the number of cells that invaded through the matrigel-coated basement membrane between the treated (losartan and candesartan) and control groups (Fig. S5E). These results were further confirmed by statistical analysis, suggesting the essential role of KLF4 in ATII-mediated GC cell metastasis. In line with the cell migration and invasion results, we also observed that losartan treatment had no significant impact on the expression of CLDN 1, 3, and 4, as well as ZO-1, in *KLF4*-deleted GC cells (Fig. S5F). These findings confirm the inability of AT1R blockers to restore TJs in GC in the absence of KLF4, thereby establishing the ATII/AT1R/KLF4 axis as a regulator of TJs in GC metastasis.

### Angiotensin II receptor type 1 blockade significantly inhibits the growth and metastasis of gastric cancer *in vivo*

The efficacy of AT1R inhibition *in vivo* was checked next. For this, GC xenografts were created using MKN45 and NCI-N87 GC cells, which were orthotopically implanted into athymic mice (Fig. 8A). Liver metastases were observed in both xenograft models, indicating the invasive and metastatic potential of the implanted GC cells (Fig. 8B). Upregulation of ATII and AT1R expressions similar to human GC was also observed in tissues collected from mouse xenografts (Fig. 8C). Interestingly, like human GC, loss of KLF4, which is otherwise expressed in normal mouse stomach, was observed in both these models (Fig. 8D). The effect of ATII inhibition *in vivo* was tested by administering AT1R antagonists, losartan (10 mg/kg) and candesartan (2 mg/kg), orally for 28 consecutive days after the tumor was detected. Results demonstrated a significant inhibition of tumor growth (as measured by wet tumor weight) and metastasis in mice treated with AT1R antagonists, losartan and candesartan, compared to untreated mice. The control MKN45 group exhibited a mean tumor weight of 375.68 ± 49.71mg (mean ± SEM). In contrast, the losartan- treated group showed a significant reduction (72%) in tumor weight (102.11 ± 24.02mg, mean ± SEM) (*p*<0.0001). A 68% reduction in tumor weight in the candesartan-treated group (117.99 ± 14.92mg, mean ± SEM) compared to the untreated group (*p*<0.0001) was also observed. Similarly, in the NCI-N87 orthotopic GC model, the untreated group had a mean tumor weight of 241.87 ± 38.92mg (mean ± SEM). The losartan treated group showed a significant 69% reduction in tumor weight (73.18 ± 11.96mg, mean ± SEM) (*p=*0.0001), and candesartan-treated group a 55% reduction (106.50 ± 15.80mg, mean ± SEM) (p=0.0019) compared to the untreated group (Fig. 8E). Importantly, liver metastasis was significantly reduced in the losartan- and candesartan-treated groups (Fig. 8F) compared to untreated groups, indicating that AT1R blockers are highly effective in restricting GC growth and metastasis in mouse models of GC.

**Fig. 8.**
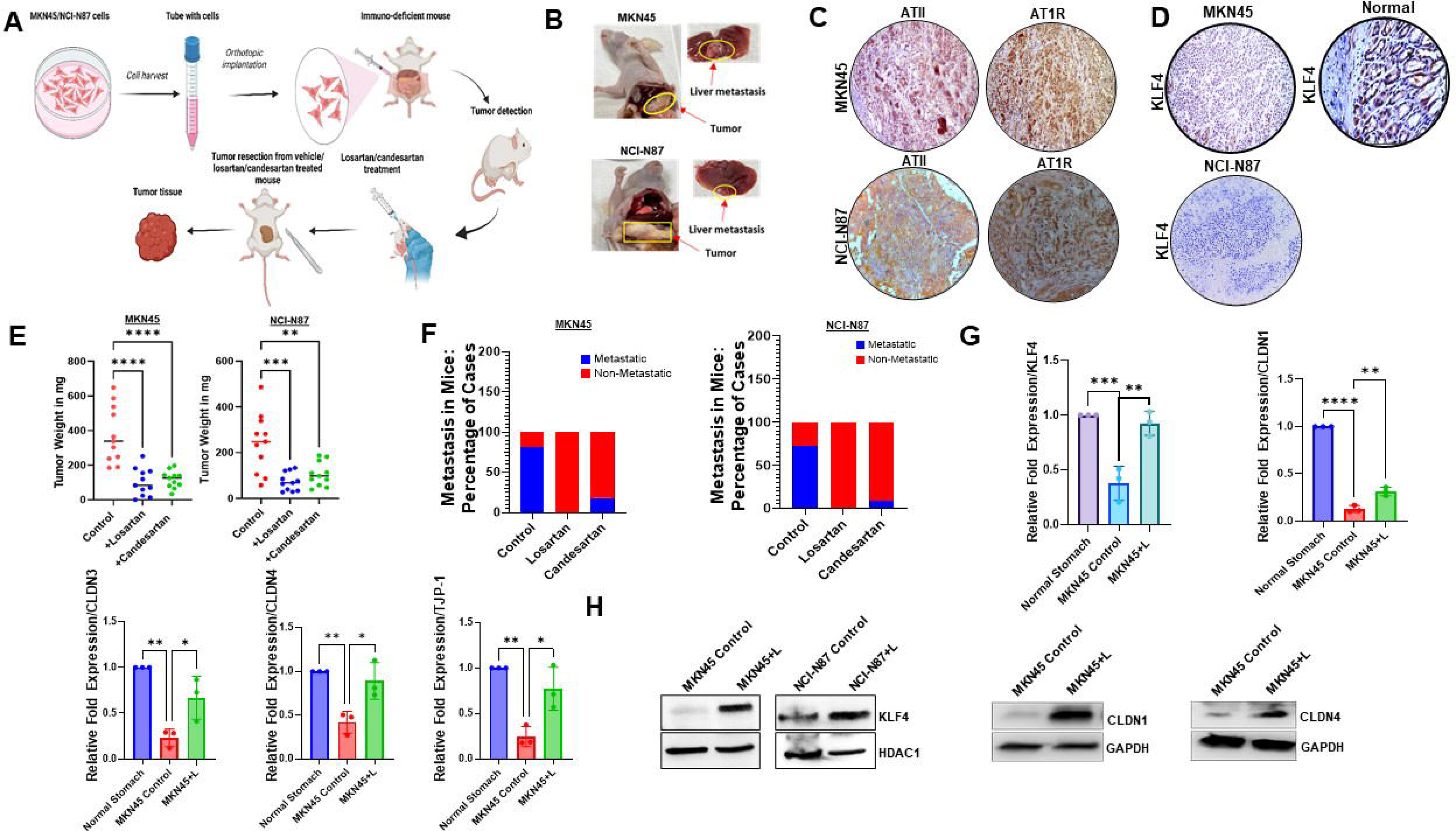
Inhibition of AT1R suppresses GC growth and metastasis *in vivo.* **(A)** Schematic representation of MKN45 and NCI-N87 GC orthotopic implantations followed by treatment with AT1R blockers in MKN45 and NCI-N87 tumor-bearing athymic mice (created by BioRender). (**B)** Representative images of orthotopic GC in mice with visible liver metastasis transplanted with MKN45 and NCI-N87 cells. (**C)** Orthotopic GC tumors show high expression of ATII and AT1R (*n =* 11 per group). Magnification, 200X (**D)** Immunohistochemistry shows a significant change in expression of KLF4 in MKN45 and NCI-N87 orthotopic tumors, which are highly metastatic to the liver. Strong KLF4 expression is observed in normal mouse stomach tissues (*n =* 11). Magnification, 200X **(E)** *In vivo* administration of AT1R blockers, losartan and candesartan, for 28 consecutive days significantly reduces tumor growth in MKN45 and NCI-N87 orthotopic xenograft models, as measured by the wet tumor weight. Each dot represents the weight of an individual mouse tumor. Data represented as mean±SEM. Significance was determined by 1-way ANOVA with Dunnett’s multiple comparisons test. **(F)** Tumor-bearing mice treated with AT1R blockers losartan and candesartan show a significant reduction in distant metastasis. **(G)** qRT-PCR analysis shows significant upregulation of *KLF4, CLDN1, CLDN3, CLDN4, and TJP1* transcripts in GC tissues from losartan-treated mice compared with untreated mice. For mRNA quantification, genes were normalized to the housekeeping gene GAPDH. Data represented as mean±SD and analyzed using 1-way ANOVA with Turkey’s multiple comparisons test (n = 3). (**H)** Western blot analysis further confirms the significant increase in the expression of KLF4, Claudin 1, and Claudin 4 in mouse GC tissues upon losartan treatment. *\*P* < 0.05*; **P* < 0.01*; ***P* < 0.001*; ****P* < 0.0001. GC, gastric cancer; IHC, immunohistochemistry; TJ, tight junction.

### Angiotensin II receptor type 1 blockade enhances tight junction protein expression in gastric cancer models

Next, we examined the TJ status in normal mouse gastric tissues and tumor xenografts. The TJ proteins were strongly expressed in the normal mouse stomach (Fig. S6A), whereas orthotopic GC tumors exhibited significant downregulation of TJ proteins, along with reduced KLF4 expression at the transcript level. Interestingly, losartan treatment significantly restored this phenotype in xenograft tumors, with increased expression of TJ genes and KLF4 compared with untreated controls (Fig. 8G). Similarly, western blot analysis showed notable upregulation of CLDN1 and CLDN4, and increased KLF4, in tumors from losartan-treated mice (Fig. 8H). These findings indicate that AT1R inhibition *in vivo* not only suppresses GC growth and metastasis but also restores TJ protein expression in gastric tumors.

## DISCUSSION

The key role of TJs within the tissue is to form intercellular barriers that maintains ion gradients across cell layers by allowing selective ions and molecules to move across the cell membrane and the paracellular space [30]. In addition, they have also garnered increasing attention for promoting adhesive interactions between adjacent cells, particularly in epithelial cells [7]. TJ abnormalities accordingly have been consistently linked to disease progression and poorer outcomes in GC, with changes in the expression and localization of several key TJ components being associated with different aspects of carcinogenesis, such as cell proliferation and loss of epithelial cell polarity, leading to migration and invasion. TJ evaluation and targeted manipulation have thus emerged as an important area of GC research, both as a window into disease biology and as a potential therapeutic strategy [31,32]. However, the complex and dynamic nature of TJs, combined with inconsistent and, in some cases, contradictory reports on individual TJ proteins in GC development and progression, has made progress difficult. Prior work has largely identified changes in expressions and localization within the TME and their effects on disease progression, but has provided little information on the molecular events underlying TJ changes or GC disruption.

In this regard, the present findings advance the field by connecting a ligand-receptor signaling pathway within the GC TME that directly compromises TJ integrity by repressing multiple TJ components in gastric epithelial cancer cells. Specifically, we uncover a previously unrecognized autocrine function of cancer cell-derived ATII and its receptor AT1R, which are overexpressed in the GC TME, in weakening the TJ barrier and facilitating the easy escape of GC cells from the primary tumor site. These findings extend prior observations from our and other laboratories on the existence and functions of local tRAS [13,33], which, unlike the systemic RAS, which primarily regulates cardiovascular homeostasis and electrolyte balance, plays an important role in tissue growth and epithelial organization. Dysregulation of tRAS has been reported in multiple malignancies, including GC, where tumor cell-derived ATII has been shown to influence cancer progression and outcome, acting primarily on endothelial cells, immune cells, and stromal fibroblasts, creating a tumor supportive niche within the TME [34–38]. Despite those reports, the molecular mechanisms by which tumor cell-derived ATII contributes to tumor progression are not fully understood, although interestingly, pharmacologic manipulation of this pathway has been effective [36].

Our study unveils an intriguing connection between ATII/AT1R overexpression and TJ dysregulation in GC, which is mediated through suppression of KLF4 or gut enriched KLF (GKLF), a prominent member of the zinc finger containing KLF transcription factor family known for its fundamental role in cell proliferation, cellular pluripotency and maintenance of epithelial cell layer and cellular integrity [20,39–46]. KLF4 has previously been implicated in cancer.

However, unlike other members of the KLF family, which either promote tumor progression or play a suppressive role, KLF4 has attracted significant attention for its context-dependent roles that vary by cancer type and can promote or suppress tumorigenesis [47,48]. In GC, reports of reduced expression indicate that KLF4 plays a suppressive role, although its precise contribution remains unclear to date. Here, we provide compelling evidence that defines the entire molecular cascade, identifies the upstream regulatory events that trigger its loss in GC cells, and characterizes the downstream molecular events that drive cell detachment and metastasis through TJ dysfunction. The link between the ATII-AT1R axis, KLF4 suppression, and reduced synthesis of key claudins (CLDN1, 3, 4) and ZO-1 in GC establishes a novel mechanistic regulatory scaffold and an actionable target to restore TJs and improve disease outcomes.

From a translational perspective, our study is particularly significant given the long-standing challenge of therapeutically manipulating TJs *in vivo*. The current study identifies an affordable, translatable therapeutic strategy that can be used in vivo to counter the molecular changes that drive metastatic progression of GC by restoring epithelial integrity. While the effects of AT1R inhibition on KLF4 and TJ restoration in GC cells and *in vivo* orthotopic xenograft models were primarily evaluated in this study, the findings provide a strong rationale for validation in larger, clinically annotated patient cohorts. Future studies should also explore GC subtype-specific dependencies on the ATII/AT1R axis, given the molecular heterogeneity of the disease. Collectively, our study identifies a novel, targetable signaling pathway that governs TJ integrity in GC and highlights restoration of TJ function as a promising therapeutic strategy to limit tumor invasion and metastasis.

## MATERIALS AND METHODS

### Human tissues

Metastatic and non-metastatic clinical specimens from male and female patients, adjacent normal/benign tissues, and normal gastric tissues were obtained from The Ohio State University tissue archive and the University of South Alabama Health biobank and histology core facility in accordance with institutional IRB guidelines. These formalin-fixed, paraffin-embedded tissue sections were used for immunohistochemical analysis.

### Cell culture

MKN45 (JCRB, Osaka, Japan) and NCI-N87 (ATCC, Manassas, VA) gastric cancer cell lines were cultured in RPMI medium, and AGS (ATCC, Manassas, VA) was cultured in F12-K medium supplemented with 10% FBS (ATCC, Manassas, VA). Cells were maintained in a 37°C humidified incubator with 5% CO_2_. All cell lines were validated by short tandem repeat DNA fingerprinting at ATCC and were found to be free of mycoplasma contamination.

### CRISPR/Cas9 KLF4 gene knockout in gastric cancer cells

The KLF4 gene knockout was performed using the KN2.0 Human Gene Knockout Kit via CRISPR non-homology mediated genome repair (Cat No: KN406691, OriGene, Rockville, USA). The gastric cancer cells (MKN45 and NCI-N87) were plated in a 6-well plate (3x10^5^ cells) and incubated at 37℃ with 5% CO_2_ to reach 60-70% confluency. 1µg of the gRNA vectors (KN406691G1, KN406691G2) was diluted in 250µL of Opti-MEM I (Gibco, Cat No: 31985070). 1µg of the linear donor DNA containing LoxP-EF1A-tGFP-P2A-Puro-LoxP (KN406691D) was added into the same 250µL of Opti-MEM I. 6µL of Turbofectin 8.0 was added into the diluted DNA, and all the components were mixed gently. The mixtures were incubated for 15 minutes at room temperature. The mixture was added dropwise to the cells, which were incubated in a 5% CO_2_ incubator. 48 hours post-transfection, the cells were passaged at a 1:10 ratio and cultured for an additional 48 hours. The cells underwent four subsequent passages under standard culture conditions, followed by puromycin (2µg/ml) selection [49], media change, and then allowed to grow for 48 hours. Once the cells had acquired adequate confluence, they were plated onto 10cm cell culture dishes. KLF4 gene knockout was confirmed with western blot and qRT-PCR. During generation, we also included scramble control gRNA (Scr) as a negative control (Cat No: KN303984, SKU GE100003, OriGene, Rockville, USA). KLF4 knockout MKN45 (KLFKOMKN45) and NCI-N87 (KLF4KONCI-N87) clones were expanded, cryopreserved, and used for further experiments.

### Mouse tumor models and treatment

All mouse experiments were performed according to protocols approved by the Institutional Animal Care and Use Committee (IACUC) at The Ohio State University, Columbus, OH, and the University of South Alabama, Mobile, AL. For the development of orthotopic MKN45 and NCI-N87 tumor models, athymic nude mice of both sexes, aged 6-8 weeks, were purchased from The Jackson Laboratory, Bar Harbor, ME, and the models were created as described previously [17,50]. All mice were housed in ventilated cages (max 4 mice/cage) under standard conditions in the institutional animal facility, with controlled temperature (22LJ±LJ2°C), humidity (55LJ±LJ10%), and a 12h light/dark cycle. Animals had a minimum 7-day acclimatization period after arrival and before any procedures were performed. A combination of ketamine and xylazine (100 mg/kg and 10 mg/kg, i.p.) was used to anesthetize the mice. MKN45 and NCI-N87 cells (1 × 10^6^ cells in 50μL 1X PBS) were transplanted to the serosal side of the stomach with the bevel facing up. The stomach was carefully returned to the abdominal cavity, and absorbable sutures were used in a 2-step closure process; first, the peritoneal layers were closed, followed by the skin incision [13,17,51,52]. Buprenorphine (2mg/kg BW) or Ethiqa (0.05ml per 20gm mouse) was administered prior to surgery and after recovery for the next 72 hours. Animals with similar sized tumors in stomach were divided randomly into control and treatment groups. Eleven mice were used per group to ensure adequate power, given a 1.7-fold effect on tumor size and metastatic load (CV=50% and alpha=0.025/2 for two two-group comparisons). None of the animals were excluded, and analysis was conducted on all these mice. None of the animals reached the humane endpoint criteria (weight loss of more than 20% and interference with the normal functioning of the vital organs and physical activities). Treatment group mice were treated with losartan (10mg/kg X 28 days, oral gavage) [53] and candesartan (2mg/kg X 28 days, Oral gavage) [54]. Mice were sacrificed on the 29^th^ day. At necropsy, tumors were harvested from the primary site, and the wet tumor weights were recorded and compared among different groups.

### Immunohistochemistry staining

After collection, all tissues were either snap-frozen or fixed in 10% neutral buffered formalin. The formalin-fixed sections were processed for paraffin sectioning. The sections were cut at a thickness of 5μm for histology or immunostaining experiments. Tissue sections were deparaffinized at 65°C for 15 minutes and hydrated through serial baths with xylene and alcohol before staining. Antigen retrieval was done in a pre-heat steamer (2100 Antigen Retriever, Electron Microscopy Sciences) using sodium citrate buffer (10 mM sodium citrate, 0.05% Tween 20, pH 6.0) for 2 hours. Slides were allowed to cool at room temperature for 20 minutes. Tissue sections were washed 5 times in 1X phosphate-buffered saline (PBS) and then immersed in methanol containing 3% H_2_O_2_ for 5 minutes at room temperature. The sections were again rinsed with PBS and incubated with 2.5% normal horse serum (Vector Laboratories, CA) at room temperature for 20 minutes to block the nonspecific binding sites and were stained for ATII (Cat# NB100-62346 Novus Biologicals, CO), AT1R (Cat# ab124505 Abcam, MA), KLF4 (Cat# ab216968 Abcam, MA and Cat# 4038S Cell Signaling Technology, Danvers, MA), CLDN1 (Cat# H00009076-M01 Abnova, Taipei, Taiwan), CLDN3 (Cat# NBP2-46299 Novus Biologicals, CO), CLDN4 (Cat# ab53156 Abcam, MA), ZO-1 (Cat# NBP2-80141 Novus Biologicals, CO). The slides were finally visualized; stained cells were counted/ scored in a blinded manner by two independent investigators and photographed using the Revolve model of ECHO: A BICO Company fluorescent microscope [17,52]. The tissue staining percentage was measured on a scale of 0-4 (0= no stain, 1 = <10% positive staining, 2 = 10–50% positive staining, 3 = 51–80% positive staining, and 4 = >80% positive staining). Staining intensity was measured on a scale of 0-3 (0 = no reaction, 1 = mild reaction, 2 = moderate reaction, and 3 = intense reaction). The score for each section was calculated by multiplying the two values obtained to normalize the heterogeneity of staining and account for the multifocal nature of the tumor [52,55].

### Immunofluorescence staining

Immunofluorescence staining was performed on MKN45, NCIlzlN87, and KLF4 knockout MKN45 gastric cancer cells. These cells were grown on FluoroDish glass-bottom plates and treated with losartan (1 µM) for 24 hours. This protocol was used to assess the expression of CLDN1 (Cat# H00009076-M01, Abnova, Taipei, Taiwan), CLDN4 (Cat# ab53156, Abcam, MA), and ZO-1 (Cat# NBP2-80141, Novus Biologicals, CO). Nuclear localization of KLF4 (Cat# ab216968, Abcam, MA; Cat# 4038S, Cell Signaling Technology, Danvers, MA) was evaluated after treating gastric cancer cells with losartan (1 µM) for 1, 2, 4, and 8 hours. Cells were fixed with 4% formaldehyde for 20 minutes at room temperature. After fixation, cells were washed 3 times with 1X PBS, then permeabilized with 0.1% Triton X-100 in 1X PBS for 20 minutes at room temperature. At the end of incubation, the solution was discarded, and the cells were washed three times with 1X PBS. Cells were then blocked with blocking buffer (1% BSA in 1X PBS) for 1 hour at room temperature. After blocking, cells were incubated with primary antibodies, diluted in 0.1% BSA in 1X PBS (CLDN1, CLDN4, ZO-1, and KLF4) at 4°C overnight. After incubation, the primary antibody was discarded, and the cells were washed three times with wash buffer. The cells were incubated with secondary antibodies, diluted in 0.1% BSA in 1X PBS, for 1 hour at room temperature in the dark. This was followed by three washes with wash buffer and a final wash with 1X PBS. Cells were mounted with Invitrogen ProLong Diamond Antifade Mountant with DAPI (Cat# P36962, Invitrogen, MA), and a coverslip was placed at the top. The stained FluoroDish was left overnight at room temperature in the dark. The samples were then visualized under a confocal scanning microscope (Nikon Eclipse Ti), analyzed, and compared across groups. Images were analyzed and processed using NIH ImageJ (Version 1.52p).

### Immunofluorescence image quantification

To quantify fluorescence intensity in stained cells, we used a custom macro script written in Nikon’s NIS-Elements software, utilizing its built-in algorithms for image processing and measurement. Briefly, maximum-projection images of the collected multi-channel confocal Z-stacks were generated, creating a single image containing the brightest value for each XY pixel location for each channel in the stack. All subsequent image processing and measurements were performed with the maximum intensity projection images. To measure staining intensity in the green channel while excluding background pixels from the intensity calculations, the green channel was first thresholded with a relatively low intensity value (165 out of a possible 4095) to generate a binary mask representing all positively stained cells. Morphological smoothing, cleaning, separation, and size restriction were then applied to the binary mask to smooth cell contours and exclude shot noise in the image. The mean intensity of the pixels included in the binary mask was then measured in arbitrary fluorescence units (A.F.U.), output to Excel, and statistically analyzed [56].

### Western blot analysis

The antibodies ATII (Cat# NB100-62346 Novus Biologicals, CO), AT1R (Cat# ab124505 Abcam, MA), KLF4 (Cat# ab216968 Abcam, MA and Cat# 4038S Cell Signaling Technology, Danvers, MA), claudin 1 (Cat# H00009076-M01 Abnova, Taipei, Taiwan), claudin 3 (Cat# NBP2-46299 Novus Biologicals, CO), claudin 4 (Cat# ab53156, Abcam, MA), Zonula occludens 1 (Cat# NBP2-80141 Novus Biologicals, CO), GAPDH (Cat#2118 Cell Signaling Technology, MA), HDAC1 (Cat# NB100-56340 Novus Biologicals, CO) were used for western blotting (Table S1). RIPA (Boston Bioproducts, MA), containing protease inhibitor cocktail, PMSF, and sodium orthovanadate (Sigma, MO), was used for lysing the cells. The proteins were estimated using the Bradford protein assay, Bio-Rad Laboratories, CA. An equal amount of (50μg of cell lysates) was electrophoresed using a polyacrylamide gel. The proteins were transferred to a 0.2-μm PVDF membrane using a Bio-Rad Trans-Blot TurboTM Transfer Pack. The membranes were blocked with 5% nonfat dried milk in TBS (5%) and then incubated with primary and HRP-conjugated secondary antibodies. A Protein molecular weight marker (Precision Plus protein Standards, Bio-Rad Laboratories, CA) was run on the same gel to determine the molecular weight. The bands were visualized with a C-Digit blot scanner [17,52]. Densitometry analysis was performed to measure the representative bands and compare the groups. GAPDH and HDAC1 served as internal controls.

### Nuclear and cytoplasmic fractionation

MKN45 and NCI-N87 (control), KLF4 knockout cells were harvested using trypsin-EDTA and centrifuged at 500×g for 5LJminutes. Cell pellets were washed with PBS and resuspended in ice-cold CER I buffer (Cat#78833, Thermo Fisher Scientific, CA) according to the manufacturer’s protocol. After 10LJminutes of incubation on ice, CER II was added, followed by brief vortexing and centrifugation at 16,000×g for 5LJminutes. The supernatant (cytoplasmic fraction) was collected. The nuclear pellet was then resuspended in ice-cold NER buffer, vortexed intermittently over 40LJminutes, and centrifuged at 16,000×g for 10LJminutes. The resulting supernatant (nuclear extract) was collected and stored at -80°C [57].

### Cell proliferation and cell growth determination

Cell proliferation was assessed using the PrestoBlue cell viability reagent (Cat# A13261, Thermo Fisher Scientific, CA). MKN45 and NCI-N87 cells were seeded in 96-well plates at 1 × 10^4^ cells/well in 100 µL complete medium and allowed to adhere overnight. The next day, cells were treated with losartan (1µM) and candesartan (1µM) and incubated at 37 °C, 5% CO₂ for 24 hours. At the end of treatment, 10 µL PrestoBlue reagent (10% v/v) was added directly to each well, including medium-only blanks, mixed gently, and incubated for 30 min at 37 °C, 5% CO₂, protected from light. Absorbance was measured at 570 nm with a 600 nm reference using a microplate reader. For determination of cell growth rate or cell doubling time, Wild-type and knockout MKN45 and NCI-N87 gastric cancer cells were seeded in 96-well plates at an initial density of 1 × 10^4^ cells/well in complete growth medium and allowed to grow for 72 hours under standard conditions (37 °C, 5% CO₂). At 72 h, viable cell numbers were determined by trypan blue exclusion using a hemocytometer. The doubling time was calculated with a formula {Duration x *ln* (2) / *ln* (Final concentration/Initial concentration)}. The growth rate of the cells was calculated by using the formula {*ln*(Final concentration/Initial concentration)/Duration} [58].

### Scratch wound assay

Cells were cultured in a 96-well plate and allowed to grow until they reached near confluence. Following that, cells were starved for 12 hours. The scratch was created with a 10µl pipette tip in one direction. Losartan (1µM) and candesartan (1µM) were added to the treated groups. After the scratch, the images were taken, and the scratch area was measured as A0. Well bottoms were marked with a fine-tip marker, which served as the reference point for further measurements. The cells were incubated at 37 °C for 24 hours. Cell movements were monitored and imaged every 2-4 hours using a phase contrast microscope. Experiments were terminated when wound gap closure was first observed in any of the experimental groups. Images were taken from the same field, and the gap was calculated in all the groups at the same time point. Wound closure was calculated by measuring the average area between edges of the two migrating columns and expressed as % wound closure [% wound closureLJ=LJ{(A0 − At/A0)LJ×LJ100} where A0 is the initial wound area between two migrating cell fronts and At is the final area between two migrating cell fronts and expressed as relative wound closure compared to the control groups [17,52,59].

### *In vitro* cell invasion assay

Cell invasion was analyzed using Transwell assays. A Corning® BioCoat™ Matrigel® Invasion Chamber with an 8.0-µm PET membrane (Corning, NY) was used to investigate cell migration [17,52,60]. MKN45 (1x10^3^) and NCI-N87 (1x10^4^) cells were added to the upper chamber containing serum-free media and pre-coated with Matrigel. The medium containing 10% FBS, which served as the chemoattractant, was added to the lower chamber. Once the cells in the upper chamber had settled, the AT1R inhibitors losartan (1µM) and candesartan (1µM) were added to the upper chamber, and the cells were allowed to invade for 24 hours at 37℃ with 5% CO2. The inserts were taken out, and cells in the upper chamber were removed using a cotton swab. The cells attached to the bottom of the membrane were fixed and stained with Hema 3™ Manual Staining System and Stat Pack (Fischer Scientific, NH). After staining, the number of cells that migrated across the coated membrane was visualized using the Revolve ECHO: A BICO fluorescent microscope. After standardization and compensation with the negative control, the number of migrated cells in the control and treatment groups was calculated and compared among groups.

### Quantitative Real-time PCR

The RNeasy Mini Kit (Qiagen, MD) was used according to the manufacturer’s protocol to isolate total RNA from MKN45 and NCI-N87 cells and from mouse tumor models. The quantity and quality of the RNA were determined with a NanoDrop spectrophotometer (Thermo Fisher Scientific, CA). cDNA synthesis was done using the RevertAid™ premium first-strand cDNA synthesis system from Thermo Scientific using oligo dT primers. Quantitative Real Time-PCR (qRT-PCR) was performed with SYBR green mix (Cat #: K0221, Thermo Fisher Scientific, CA) using QuantStudio ™ 3 Real-Time PCR System (Applied Biosystem, MA) to determine expressions of *KLF4, AGT, AGTR1, CLDN1, CLDN3, CLDN4, AND TJP-1* in the samples with first-strand cDNA as templates. Gene sequences were identified and corresponding PCR primers designed using the Genebank database. GAPDH was used as an endogenous control. Arbitrary units were used to express the relative expression, and the fold change in gene expression was determined by the ΔCt method. Primers used: *KLF4* (forward 5′-CATCTCAAGGCACACCTGCGAA-3′, reverse 5′-TCGGTCGCATTTTTGGCACTGG -3′); *CLDN1* (forward 5′-GTCTTTGACTCCTTGCTGAATCTG-3′, reverse 5′-CACCTCATCGTCTTCCAAGCAC -3′); *CLDN3* (forward 5′-GCCTTCATCGGCAGCAACATCA-3′, reverse 5′-AGCGAGTCGTTACACCTTGCACT-3′); *CLDN4* (forward 5′-AGTGCAAGGTGTACGACTCGCT-3′, reverse 5′-CGCTTTCATCCTCCAGGCAGTT-3′); *TJP-1* (forward 5′-GTCCAGAATCTCGGAAAAGTGCC -3′, reverse 5′-CTTTCAGCGCACCATACCAACC-3′); *AGT* (forward 5’-TGGACAGCACCCTGGCTTTCAA - 3′, reverse 5′-ACACTGAGGTGCTGTTGTCCAC-3′); *AGTR1* (forward 5’-CAGCGTCAGTTTCAACCTGTACG - 3′, reverse 5′-GCAGGTGACTTTGGCTACAAGC-3′) and *GAPDH* (forward 5’-GTCTCCTCTGACTTCAACAGCG- 3′, reverse 5′-ACCACCCTGTTGCTGTAGCCAA-3′) (Invitrogen, CA). Primer sequences for gene expression studies are provided in Table S2.

### Chromatin Immunoprecipitation (ChIP) Assay

The genomic database GeneCopoeia was used to identify transcription start sites for genes encoding key TJ proteins. For each gene, upstream sequences flanking the TSS (typically from - 1000 to +100 base pairs) known to harbor promoter elements and potential transcription factor binding sites were extracted. The JASPAR database was utilized to analyze the extracted promoter sequences, with a focus on the KLF4 binding motif. The binding sites with higher relative scores, indicating a stronger likelihood of KLF4 binding, were selected and investigated. Following the manufacturer’s protocol, ChIP assays were performed using the SimpleChIP® Enzymatic Chromatin IP Kit (Cell Signaling Technology, USA; Cat # 9003S) [61]. MKN45 cells (3x10^5^) were cultured in 100mm dishes to 50-60% confluence and treated with losartan at 1µM for 24 hours. After that, the cells were cross-linked using 1% formaldehyde (Sigma-Aldrich, Cat # 252549) to preserve protein-DNA interactions. Cells were then lysed, and chromatin was digested using Micrococcal Nuclease (MNase) (Cell Signaling Technology, USA; Cat # 10011) per the kit instructions. Following digestion, chromatin was immunoprecipitated using an anti-KLF4 antibody. For control experiments, IgG (Cell Signaling Technology, USA; Cat # 2729) was used as a negative control, and Histone H3 (Cell Signaling Technology, USA; Cat # 4620) was used as a positive control. Immunoprecipitated chromatin complexes were eluted, and DNA was recovered and purified following the manufacturer’s instructions. The purified DNA was subjected to qRT-PCR to detect the binding of KLF4 to the promoter regions of *CLDN1*, *CLDN3*, *CLDN4,* and *TJP1* using promoter sequence-specific primer sets. The fold change in ChIP DNA was determined using qRT-PCR. Primer sequences for the ChIP assay are provided in Table S3.

### RNA isolation from cell lines

RNA was extracted from MKN45 (control), KLF4 knockout, and losartan-treated cells using Qiagen’s RNeasy MINI Kit per the manufacturer’s instructions. The isolated RNA was quantified and qualitatively assessed using a Qubit Fluorometer (Thermo Fisher Scientific) and a Bioanalyzer (Agilent2100), according to the manufacturer’s instructions. The total RNA, with a concentration of 20 ng/µl, an RNA integrity score ≥4.0, and an OD260/280 ratio ≥2.0, was then sent to Novogene (https://en.novogene.com/) for mRNA sequencing (WBI).

### Library Construction, Quality Control, and Sequencing

Messenger RNA was purified from total RNA using poly-T oligo-attached magnetic beads. The first strand cDNA was synthesized after fragmentation by using random hexamer primers. This was followed by the second strand cDNA synthesis using dTTP for a non-directional library. The library was ready after end repair, A-tailing, adapter ligation, size selection, amplification, and purification. The library was checked using Qubit, and real-time PCR was employed for quantification. A bioanalyzer was used to determine the size distribution. Illumina platforms were used to sequence pooled quantified libraries based on effective library concentration and data volume [62,63].

### RNA-sequencing analysis

RNA sequencing was performed in duplicate on the Illumina NovaSeq 6000 using paired-end sequencing (150 bp). There were ∼40 million reads for each sample. The processed FASTQ files were mapped to the reference genome using HISAT2, and feature counts were used for gene expression quantification. The raw counts were normalized and expressed as Fragments Per Kilobase of transcript per Million mapped reads (FPKM) to estimate the gene expression levels and enable comparison across samples. For the differential gene expression analysis, the DESeq2 R package (version 1.20.1) was used to compare controls to losartan and knockout samples, as previously described [64]. Gene set enrichment analysis (preranked GSEA) was carried out in R (version 4.5.1) using the ranked DESeq2 statistics and gene sets downloaded from the GSEA/MsigDB resource [65]. DESeq2 Wald statistics were ranked from upregulated to downregulated and used as input for preranked gene set enrichment analysis (GSEA) using clusterProfiler, enrichplot, and fgsea, and the Benjamini-Hochberg procedure was applied to control false discovery rate (FDR) to obtain enrichment score, NES, nominal P, and FDR for the TJ pathway. The gene ontology and pathways of differentially expressed genes (DEGs) were analyzed with BigOmics Analytics [24]. The volcano plot and heatmaps of differentially expressed genes were generated using the SRplot online tool [66].

### TCGA analysis

The TCGA data on *AGT* and *AGTR1* expressions were obtained from the TCGA data portal (http://ualcan.path.uab.edu) [67,68]. The data were accessed in January 2026, and RNAseq data were available for a total of 415 primary tumors and 34 normal samples. Gene expression data is represented as normalized RNA-seq counts (Transcripts Per Million, TPM) from the TCGA dataset. The expression levels of *AGT* and *AGTR1* were analyzed in tumor tissues compared to adjacent normal stomach. Also, the data were stratified by tumor stage at different stages for subgroup analysis (i.e., Stage I, II, III, and IV). To compare the impact of AGT and AGTR1 expression on overall survival (OS) and disease-free survival (DFS) in stomach cancer patients, survival analysis of *AGT* and *AGTR1* for stomach adenocarcinoma patients in the TCGA-STAD profile was compared in the Gene Expression Profiling Interactive Analysis web service (GEPIA, http://gepia.cancer-pku.cn/index.html) [69]. The OS analysis was performed using the Kaplan-Meier survival curve provided by GEPIA. We plotted the survival curve and extracted information on median survival time, hazard ratio (HR), and the log-rank *p-*value to determine the statistical significance of survival differences between the high and low expression groups. Similarly, DFS analysis was performed using GEPIA’s survival function. The same statistical parameters (median survival time, HR, and log-rank *p-*value) were used to assess the influence of *AGT* and *AGTR1* expression on DFS. A log-rank test was used to analyze the statistical significance of survival in high and low *AGT* and *AGTR1* expression groups. Statistical significance was set at *p <* 0.05.

### TCGA**fll**STAD transcriptomic analysis of AGT/AGTR1 and tight junction genes

The gene expression data were extracted from The Cancer Genome Atlas Stomach Adenocarcinoma (TCGA-STAD) cohort via the NCI Genomic Data Commons (GDC) portal (gene model: GENCODE v36). The RNA-sequencing data were retrieved as TPM (Transcripts Per Million) values and log2-transformed [log2(TPM+1)] before analysis. The study analyzed n=200 tumor samples with available expression of *AGT, AGTR1,* and 30 TJ related genes. The samples were stratified for *AGT* and *AGTR1* expression using a median split, with the bottom 50% (n=100) designated as the low *AGT* and *AGTR1* expression group and the top 50% (n=100) as the high *AGT* and *AGTR1* expression group. The Median log₂ (TPM + 1) cutoff values used were 7.967 for *AGT* and 6.940 for *AGTR1*. The differences in the expression of TJ genes between the low and high *AGT* and *AGTR1* groups were assessed using the Wilcoxon rank-sum test (Mann-Whitney U, two-sided), a non-parametric test. A *p -*value *< 0.05* was set as statistical significance. All analyses were performed in R version 4.5.1.

### Statistics

All statistical analyses were conducted in a blinded manner. GraphPad Prism 8.1.0 (San Diego, California, USA) was used to analyze quantitative data from *in vivo* and *in vitro* experimental studies. A two-sample t-test was used to analyze data from experiments involving two groups. Data from studies with multiple experimental groups were analyzed using ANOVA. Quantitative data are presented as mean ± SD. For correlation studies, the correlation between two variables was analyzed using Spearman’s correlation method. Each RNA-seq experiment was performed in two independent biological replicates. For RNA-seq, DEGs were defined as those with a fold change (FC) > 1.5 and a *p*-value <0.05. Significance levels are indicated as \**p*<0.05, *\*\*p*<0.01, *\*\*\*p*<0.001, \**\*\*\*p*<0.0001 [17,52].

## Author’s contribution statement

Conception: DC; design: DC; development of methodology: SK, PS and SG; experiment and data generation: SK, PS, SG, BK and KB; analysis and interpretation of data: DC, CS, SK, PS, SG, BK, WF, SB, EH, VRA, JA and PP; writing original draft: SK, CS and DC; writing/reviewing and editing: SK, MH, VRA, CS and DC; administrative and study supervision: DC. All authors read and approved the final manuscript.

## Ethics approval and consent to participate

All mouse experiments were performed according to protocols approved by the Institutional Animal Care and Use Committee (IACUC) at The Ohio State University, Columbus, OH, and the University of South Alabama, Mobile, AL.

## Consent for publication

All authors give consent to publish and agree on the final manuscript submission.

## Availability of data and materials

All data associated with this study are present in the paper or the Supplementary Materials. The raw and analyzed datasets generated during the study are available for research purposes from the corresponding author on reasonable request.

## Funding

This study is supported by funding from DOD, USA [W81XWH-20-1-0618 to DC]. DC also acknowledges funding received from the Mitchell Cancer Institute, the University of South Alabama, and the Robert E. Reed Gastrointestinal Oncology Research Foundation (to DC and CS).

## Declaration of competing interest

The authors declare that they have no known competing financial interests or personal relationships that could have influenced the work reported in this paper.

## Acknowledgments

We acknowledge the support of the veterinarian and vivarium staff at OSU, as well as Dr. Michele Schuler and Ms. Leigh Ann Wiggins of the USA, for our animal studies. We also acknowledge the support of USA Health Biobank, Histology, and confocal microscopy core facilities for this study.

## Supplemental Material

Fig. S1 displays tight junction gene expression associated with AGT and AGTR1 status in TCGA-STAD. Fig. S2 shows the expression of KLF4 in GC tissues compared with normal stomach as revealed by TCGA transcriptomic data and Human Protein Atlas. Fig. S3 demonstrates the expression of KLF4 in different GC cells and migratory and invasive behavior of KLF4 expression modulated GC cells. Fig. S4 reveals the negative correlation of ATII and AT1R with KLF4 by IHC and cBioPortal data. Fig. S5 shows the effect of AT1R inhibition in migratory and invasive behavior of *KLF4*- deleted GC cells. Fig. S6 provides western blot data showing baseline tight junction protein expression in normal mouse stomach tissue. Table S1 shows the list of antibodies used for immunohistochemistry, western-blot and immunofluorescence analyses. Table S2 shows the sequences of primers used for qRT-PCR. Table S3 shows the sequences of primers used in ChIP assay.

---

**Fig. S1.** Tight junction gene expressions and their association with *AGT* and *AGTR1* in TCGA-STAD. **(A)** The heatmaps display Z-score-scaled expression of TJ-related genes across TCGA-STAD tumors (n = 100 per group for each gene). The column represents individual tumors, and rows represent the TJ-related genes. The Z score blue, white, and red denotes the relative under- to over-expression across the cohort. The top bars indicate the continuous *AGT* or *AGTR1* expression and their status (Low vs High). Low *AGT* tumors generally have higher TJ gene expression, while high *AGT tumors* show downregulation of these TJ genes. AGT, angiotensin II; AGTR1, angiotensin II receptor type 1.

**Fig. S2.** KLF4 downregulation in GC is associated with poor patient prognosis. (**A)** Heatmap generated using TCGA transcriptomic data shows the expression of Kruppel-like factor (*KLF*) family genes in normal gastric (*n* = 34) and primary stomach adenocarcinoma tissues (STAD) (*n* = 415). Data indicate a consistent change in *KLF4* (downregulation or loss) and *KLF5* (upregulation) expression in human GC tissues (**B)** Box plots show significantly decreased *KLF4* and increased *KLF5* transcript levels in STAD tumors compared to normal tissues (data from TCGA). It also shows a stage-wise decrease in KLF4 expression in STAD. (**C)** Representative IHC images from The Human Protein Atlas reveal a significant loss of KLF4 in malignant gastric cancer tissue. GC, gastric cancer; STAD, stomach adenocarcinoma; TCGA, The Cancer Genome Atlas; IHC, immunohistochemistry.

**Fig. S3.** Loss of KLF4 promotes cell proliferation and metastasis of GC cells. **(A)** Representative qRT-PCR and western blot images show KLF4 expression in different GC cell lines. Data represented as mean±SD. (**B and C)** Western blot and qRT-PCR analysis confirm the deletion of KLF4 in MKN45 and NCI-N87 cells after CRISPR/Cas9-mediated knockout. For mRNA quantification, KLF4 was normalized to the housekeeping gene GAPDH. Densitometric quantification of KLF4 protein levels normalized to GAPDH from the western blots. Significance was determined by 1-way ANOVA with Dunnett’s multiple comparisons test and 2-tailed *t* test (*n* = 3) (**D).** A significant decrease in cell doubling time and an increase in cell proliferation are observed in KLF4KOMKN45 and KLF4KONCI-N87 cells compared to their scrambled control cells (Scr). Significance was determined by a 2-tailed *t* test *(n = 3).* (**E)** KLF4 knockout MKN45 and NCI-N87 cells (KLF4KOMKN45 and KLF4KONCI-N87) show markedly enhanced cell migration *in vitro*, as demonstrated by wound healing assays at 0 h and 24 h. Magnification, 40X. Transwell invasion assays revealed significantly increased invasive capacity in *KLF4*-deleted cells compared to scrambled cells. Magnification, 200X. Quantification of migrated and invaded cells is shown in bar graphs (right panels). Data represented as mean±SEM. Significance was determined by a 2-tailed t-test (migration, *n* = 3; invasion, *n* =9). *\*P <* 0.05*; **P* < 0.01*; ***P* < 0.001*;****P* < 0.0001. GC, gastric cancer; Scr, scrambled control.

**Fig. S4.** ATII/AT1R expression is negatively associated with expression KLF4 in GC. **(A)** The quantitative correlation analysis shows a significant negative correlation between the average IHC score for ATII/AT1R expression and the number of KLF4-positive cells. Correlation was assessed using Pearson’s correlation test; results are displayed in the graph (*n =* 46). **(B)** cBioPortal data show that elevated mRNA expression of *AGT* and *AGTR1* is negatively correlated with *KLF4* expression in GC. *\*P* < 0.05*; **P* < 0.01*; ***P <* 0.001*; ****P* < 0.0001. GC, gastric cancer; TJ, tight junction; AGT, angiotensin II; AGTR1, angiotensin II receptor type 1.

**Fig. S5.** AT1R inhibition fails to restore the expression of TJ proteins or inhibit the metastatic capacity of KLF4-knockout GC cells. (A-D) Wound closure and trans well assays show that neither losartan nor candesartan treatment significantly alters the migratory or invasive potential of KLF4KOMKN45 and KLF4KONCI-N87 cells compared to untreated cells (migration, Magnification 40X; invasion, Magnification 200X). **(E)** The percentage of wound closure and cell-invasive index after losartan or candesartan treatment is shown as bar diagrams (data as mean±SEM), with significance assessed by 2-tailed t-test (*n* = 12 for losartan wound closure, *n* = 9 for losartan invasion, *n* = 4 for candesartan wound closure, *n* = 5 for candesartan invasion). **(F)** Western blot analysis shows no significant change in levels of TJ proteins Claudin 1, 3, 4, and Zonula occludens 1 in losartan-treated cells. *ns*: no significance. GC, gastric cancer; TJs, tight junctions.

**Fig. S6.** Western blot analysis of TJ protein expression in normal mouse stomach tissue. **(A)** Representative western blot data show the expression levels of Claudin 1, 3, 4, and Zonula occludens 1 in normal mouse stomach tissue. GAPDH was used as a loading control. TJ, tight junction; NS, normal stomach.

## Notes

### Competing Interest Statement

The authors have declared no competing interest.

### Summary of Updates

The results section has been updated, and an additional supplementary figure (Figure S1A) has been included. Figure 2 and Figure S5 have been revised with updated data. The method section and figure legends have been modified accordingly to reflect these changes. The discussion has also been updated.

## REFERENCES AND NOTES

[1] J.Y. Park, D. Georges, C.J. Alberts, F. Bray, G. Clifford, I. Baussano, Global lifetime estimates of expected and preventable gastric cancers across 185 countries, Nat Med (2025). 10.1038/s41591-025-03793-6.

[2] R.E. Sexton, M.N. Al Hallak, M. Diab, A.S. Azmi, Gastric cancer: a comprehensive review of current and future treatment strategies, Cancer Metastasis Rev 39 (2020) 1179–1203. 10.1007/s10555-020-09925-3.

[3] J. Taieb, J. Bennouna, F. Penault-Llorca, D. Basile, E. Samalin, A. Zaanan, Treatment of gastric adenocarcinoma: A rapidly evolving landscape, Eur J Cancer 195 (2023) 113370. 10.1016/j.ejca.2023.113370.

[4] X. Sun, Y. Zhou, S. Sun, S. Qiu, M. Peng, H. Gong, J. Guo, C. Wen, Y. Zhang, Y. Xie, H. Li, L. Liang, G. Luo, W. Wu, J. Liu, W. Tan, M. Ye, Cancer cells sense solid stress to enhance metastasis by CKAP4 phase separation-mediated microtubule branching, Cell Discov 10 (2024) 114. 10.1038/s41421-024-00737-1.

[5] Y. Huang, W. Hong, X. Wei, The molecular mechanisms and therapeutic strategies of EMT in tumor progression and metastasis, J Hematol Oncol 15 (2022) 129. 10.1186/s13045-022-01347-8.

[6] P. Dhawan, A.B. Singh, N.G. Deane, Y. No, S.-R. Shiou, C. Schmidt, J. Neff, M.K. Washington, R.D. Beauchamp, Claudin-1 regulates cellular transformation and metastatic behavior in colon cancer, J Clin Invest 115 (2005) 1765–1776. 10.1172/JCI24543.

[7] T.P. Nguyen, T. Otani, M. Tsutsumi, N. Kinoshita, S. Fujiwara, T. Nemoto, T. Fujimori, M. Furuse, Tight junction membrane proteins regulate the mechanical resistance of the apical junctional complex, J Cell Biol 223 (2024) e202307104. 10.1083/jcb.202307104.

[8] J. Huang, J. Li, Y. Qu, J. Zhang, L. Zhang, X. Chen, B. Liu, Z. Zhu, The expression of claudin 1 correlates with β-catenin and is a prognostic factor of poor outcome in gastric cancer, Int J Oncol 44 (2014) 1293–1301. 10.3892/ijo.2014.2298.

[9] T.-L. Hwang, T.-T. Changchien, C.-C. Wang, C.-M. Wu, Claudin-4 expression in gastric cancer cells enhances the invasion and is associated with the increased level of matrix metalloproteinase-2 and -9 expression, Oncol Lett 8 (2014) 1367–1371. 10.3892/ol.2014.2295.

[10] S. Ohtani, M. Terashima, J. Satoh, N. Soeta, Z. Saze, S. Kashimura, F. Ohsuka, Y. Hoshino, M. Kogure, M. Gotoh, Expression of tight-junction-associated proteins in human gastric cancer: downregulation of claudin-4 correlates with tumor aggressiveness and survival, Gastric Cancer 12 (2009) 43–51. 10.1007/s10120-008-0497-0.

[11] S.J. Hagen, L.-H. Ang, Y. Zheng, S.N. Karahan, J. Wu, Y.E. Wang, T.J. Caron, A.P. Gad, S. Muthupalani, J.G. Fox, Loss of Tight Junction Protein Claudin 18 Promotes Progressive Neoplasia Development in Mouse Stomach, Gastroenterology 155 (2018) 1852–1867. 10.1053/j.gastro.2018.08.041.

[12] K.-H. Jun, J.-H. Kim, J.-H. Jung, H.-J. Choi, H.-M. Chin, Expression of claudin-7 and loss of claudin-18 correlate with poor prognosis in gastric cancer, Int J Surg 12 (2014) 156–162. 10.1016/j.ijsu.2013.11.022.

[13] C. Sarkar, R.K. Ganju, V.J. Pompili, D. Chakroborty, Enhanced peripheral dopamine impairs post-ischemic healing by suppressing angiotensin receptor type 1 expression in endothelial cells and inhibiting angiogenesis, Angiogenesis 20 (2017) 97–107. 10.1007/s10456-016-9531-8.

[14] M. Kosugi, A. Miyajima, E. Kikuchi, Y. Horiguchi, M. Murai, Angiotensin II type 1 receptor antagonist candesartan as an angiogenic inhibitor in a xenograft model of bladder cancer, Clin Cancer Res 12 (2006) 2888–2893. 10.1158/1078-0432.CCR-05-2213.

[15] A.M. Ghaleb, M.O. Nandan, S. Chanchevalap, W.B. Dalton, I.M. Hisamuddin, V.W. Yang, Krüppel-like factors 4 and 5: the yin and yang regulators of cellular proliferation, Cell Res 15 (2005) 92–96. 10.1038/sj.cr.7290271.

[16] P. Sangwung, G. Zhou, L. Nayak, E.R. Chan, S. Kumar, D.-W. Kang, R. Zhang, X. Liao, Y. Lu, K. Sugi, H. Fujioka, H. Shi, S.D. Lapping, C.C. Ghosh, S.J. Higgins, S.M. Parikh, H. Jo, M.K. Jain, KLF2 and KLF4 control endothelial identity and vascular integrity, JCI Insight 2 (2017) e91700. 10.1172/jci.insight.91700.

[17] D. Chakroborty, C. Sarkar, H. Yu, J. Wang, Z. Liu, P.S. Dasgupta, S. Basu, Dopamine stabilizes tumor blood vessels by up-regulating angiopoietin 1 expression in pericytes and Kruppel-like factor-2 expression in tumor endothelial cells, Proc Natl Acad Sci U S A 108 (2011) 20730–20735. 10.1073/pnas.1108696108.

[18] P. Ma, Y. Pan, F. Yang, Y. Fang, W. Liu, C. Zhao, T. Yu, M. Xie, X. Jing, X. Wu, C. Sun, W. Li, T. Xu, Y. Shu, KLF5-Modulated lncRNA NEAT1 Contributes to Tumorigenesis by Acting as a Scaffold for BRG1 to Silence GADD45A in Gastric Cancer, Mol Ther Nucleic Acids 22 (2020) 382–395. 10.1016/j.omtn.2020.09.003.

[19] Z. He, J. He, K. Xie, KLF4 transcription factor in tumorigenesis, Cell Death Discov 9 (2023) 118. 10.1038/s41420-023-01416-y.

[20] J.M. Shields, R.J. Christy, V.W. Yang, Identification and characterization of a gene encoding a gut-enriched Krüppel-like factor expressed during growth arrest, J Biol Chem 271 (1996) 20009–20017. 10.1074/jbc.271.33.20009.

[21] M. Shao, G.-Z. Ge, W.-J. Liu, J. Xiao, H.-J. Xia, Y. Fan, F. Zhao, B.-L. He, C. Chen, Characterization and phylogenetic analysis of Krüppel-like transcription factor (KLF) gene family in tree shrews (Tupaia belangeri chinensis), Oncotarget 8 (2017) 16325–16339. 10.18632/oncotarget.13883.

[22] M. Uhlén, L. Fagerberg, B.M. Hallström, C. Lindskog, P. Oksvold, A. Mardinoglu, Å. Sivertsson, C. Kampf, E. Sjöstedt, A. Asplund, I. Olsson, K. Edlund, E. Lundberg, S. Navani, C.A.-K. Szigyarto, J. Odeberg, D. Djureinovic, J.O. Takanen, S. Hober, T. Alm, P.-H. Edqvist, H. Berling, H. Tegel, J. Mulder, J. Rockberg, P. Nilsson, J.M. Schwenk, M. Hamsten, K. von Feilitzen, M. Forsberg, L. Persson, F. Johansson, M. Zwahlen, G. von Heijne, J. Nielsen, F. Pontén, Proteomics. Tissue-based map of the human proteome, Science 347 (2015) 1260419. 10.1126/science.1260419.

[23] F. Kong, T. Sun, X. Kong, D. Xie, Z. Li, K. Xie, Krüppel-like Factor 4 Suppresses Serine/Threonine Kinase 33 Activation and Metastasis of Gastric Cancer through Reversing Epithelial-Mesenchymal Transition, Clin Cancer Res 24 (2018) 2440–2451. 10.1158/1078-0432.CCR-17-3346.

[24] M. Akhmedov, A. Martinelli, R. Geiger, I. Kwee, Omics Playground: a comprehensive self-service platform for visualization, analytics and exploration of Big Omics Data, NAR Genom Bioinform 2 (2020) lqz019. 10.1093/nargab/lqz019.

[25] A. Tiwari, C.L. Loughner, S. Swamynathan, S.K. Swamynathan, KLF4 Plays an Essential Role in Corneal Epithelial Homeostasis by Promoting Epithelial Cell Fate and Suppressing Epithelial-Mesenchymal Transition, Invest Ophthalmol Vis Sci 58 (2017) 2785–2795. 10.1167/iovs.17-21826.

[26] G.L. Sen, L.D. Boxer, D.E. Webster, R.T. Bussat, K. Qu, B.J. Zarnegar, D. Johnston, Z. Siprashvili, P.A. Khavari, ZNF750 is a p63 target gene that induces KLF4 to drive terminal epidermal differentiation, Dev Cell 22 (2012) 669–677. 10.1016/j.devcel.2011.12.001.

[27] K.-G. Shyu, W.-P. Cheng, B.-W. Wang, Angiotensin II Downregulates MicroRNA-145 to Regulate Kruppel-like Factor 4 and Myocardin Expression in Human Coronary Arterial Smooth Muscle Cells under High Glucose Conditions, Mol Med 21 (2015) 616–625. 10.2119/molmed.2015.00041.

[28] S.J. Forrester, G.W. Booz, C.D. Sigmund, T.M. Coffman, T. Kawai, V. Rizzo, R. Scalia, S. Eguchi, Angiotensin II Signal Transduction: An Update on Mechanisms of Physiology and Pathophysiology, Physiol Rev 98 (2018) 1627–1738. 10.1152/physrev.00038.2017.

[29] I. de Bruijn, R. Kundra, B. Mastrogiacomo, T.N. Tran, L. Sikina, T. Mazor, X. Li, A. Ochoa, G. Zhao, B. Lai, A. Abeshouse, D. Baiceanu, E. Ciftci, U. Dogrusoz, A. Dufilie, Z. Erkoc, E. Garcia Lara, Z. Fu, B. Gross, C. Haynes, A. Heath, D. Higgins, P. Jagannathan, K. Kalletla, P. Kumari, J. Lindsay, A. Lisman, B. Leenknegt, P. Lukasse, D. Madela, R. Madupuri, P. van Nierop, O. Plantalech, J. Quach, A.C. Resnick, S.Y.A. Rodenburg, B.A. Satravada, F. Schaeffer, R. Sheridan, J. Singh, R. Sirohi, S.O. Sumer, S. van Hagen, A. Wang, M. Wilson, H. Zhang, K. Zhu, N. Rusk, S. Brown, J.A. Lavery, K.S. Panageas, J.E. Rudolph, M.L. LeNoue-Newton, J.L. Warner, X. Guo, H. Hunter-Zinck, T.V. Yu, S. Pilai, C. Nichols, S.M. Gardos, J. Philip, AACR Project GENIE BPC Core Team, AACR Project GENIE Consortium, K.L. Kehl, G.J. Riely, D. Schrag, J. Lee, M.V. Fiandalo, S.M. Sweeney, T.J. Pugh, C. Sander, E. Cerami, J. Gao, N. Schultz, Analysis and Visualization of Longitudinal Genomic and Clinical Data from the AACR Project GENIE Biopharma Collaborative in cBioPortal, Cancer Res 83 (2023) 3861–3867. 10.1158/0008-5472.CAN-23-0816.

[30] S. Tsukita, M. Furuse, Occludin and claudins in tight-junction strands: leading or supporting players?, Trends Cell Biol 9 (1999) 268–273. 10.1016/s0962-8924(99)01578-0.

[31] G. Grizzi, K. Venetis, N. Denaro, M. Bonomi, A. Celotti, A. Pagkali, J.C. Hahne, G. Tomasello, F. Petrelli, N. Fusco, M. Ghidini, Anti-Claudin Treatments in Gastroesophageal Adenocarcinoma: Mainstream and Upcoming Strategies, J Clin Med 12 (2023) 2973. 10.3390/jcm12082973.

[32] D. Luo, Y. Liu, Z. Lu, L. Huang, Targeted therapy and immunotherapy for gastric cancer: rational strategies, novel advancements, challenges, and future perspectives, Mol Med 31 (2025) 52. 10.1186/s10020-025-01075-y.

[33] S.J. Delforce, E.R. Lumbers, C. Corbisier de Meaultsart, Y. Wang, A. Proietto, G. Otton, J. Scurry, N.M. Verrills, R.J. Scott, K.G. Pringle, Expression of renin-angiotensin system (RAS) components in endometrial cancer, Endocr Connect 6 (2017) 9–19. 10.1530/EC-16-0082.

[34] D.R. Butcher, C.N. Parris, S.J. Crichton, F.C. Dempsey, H.N. Al-Ali, Unlocking the potential of targeting the angiotensin II type 1 receptor in cancer, Oncogene 45 (2026) 479–490. 10.1038/s41388-025-03666-9.

[35] J. Kinoshita, S. Fushida, S. Harada, Y. Yagi, H. Fujita, S. Kinami, I. Ninomiya, T. Fujimura, M. Kayahara, M. Yashiro, K. Hirakawa, T. Ohta, Local angiotensin II-generation in human gastric cancer: correlation with tumor progression through the activation of ERK1/2, NF-kappaB and survivin, Int J Oncol 34 (2009) 1573–1582. 10.3892/ijo_00000287.

[36] W. Huang, Y.-L. Wu, J. Zhong, F.-X. Jiang, X.-L. Tian, L.-F. Yu, Angiotensin II type 1 receptor antagonist suppress angiogenesis and growth of gastric cancer xenografts, Dig Dis Sci 53 (2008) 1206–1210. 10.1007/s10620-007-0009-9.

[37] G. Xie, T. Cheng, J. Lin, L. Zhang, J. Zheng, Y. Liu, G. Xie, B. Wang, Y. Yuan, Local angiotensin II contributes to tumor resistance to checkpoint immunotherapy, J Immunother Cancer 6 (2018) 88. 10.1186/s40425-018-0401-3.

[38] T. Takiguchi, F. Takahashi-Yanaga, S. Ishikane, F. Tetsuo, H. Hosoda, M. Arioka, T. Kitazono, T. Sasaguri, Angiotensin II promotes primary tumor growth and metastasis formation of murine TNBC 4T1 cells through the fibroblasts around cancer cells, Eur J Pharmacol 909 (2021) 174415. 10.1016/j.ejphar.2021.174415.

[39] T. Xiang, C. Yang, Z. Deng, D. Sun, F. Luo, Y. Chen, Krüppel-like factors family in health and disease, MedComm (2020) 5 (2024) e723. 10.1002/mco2.723.

[40] T. Suzuki, D. Sawaki, K. Aizawa, Y. Munemasa, T. Matsumura, J. Ishida, R. Nagai, Kruppel-like factor 5 shows proliferation-specific roles in vascular remodeling, direct stimulation of cell growth, and inhibition of apoptosis, J Biol Chem 284 (2009) 9549–9557. 10.1074/jbc.M806230200.

[41] C.-K. Kim, P. He, A.B. Bialkowska, V.W. Yang, SP and KLF Transcription Factors in Digestive Physiology and Diseases, Gastroenterology 152 (2017) 1845–1875. 10.1053/j.gastro.2017.03.035.

[42] Y. Gao, Q. Cao, L. Lu, X. Zhang, Z. Zhang, X. Dong, W. Jia, Y. Cao, Kruppel-like factor family genes are expressed during Xenopus embryogenesis and involved in germ layer formation and body axis patterning, Dev Dyn 244 (2015) 1328–1346. 10.1002/dvdy.24310.

[43] G. Ito, M. Uchiyama, M. Kondo, S. Mori, N. Usami, O. Maeda, T. Kawabe, Y. Hasegawa, K. Shimokata, Y. Sekido, Krüppel-like factor 6 is frequently down-regulated and induces apoptosis in non-small cell lung cancer cells, Cancer Res 64 (2004) 3838–3843. 10.1158/0008-5472.CAN-04-0185.

[44] H.L. Reeves, G. Narla, O. Ogunbiyi, A.I. Haq, A. Katz, S. Benzeno, E. Hod, N. Harpaz, S. Goldberg, S. Tal-Kremer, F.J. Eng, M.J.P. Arthur, J.A. Martignetti, S.L. Friedman, Kruppel-like factor 6 (KLF6) is a tumor-suppressor gene frequently inactivated in colorectal cancer, Gastroenterology 126 (2004) 1090–1103. 10.1053/j.gastro.2004.01.005.

[45] J. Lin, H. Tan, Y. Nie, D. Wu, W. Zheng, W. Lin, Z. Zhu, B. Yang, X. Chen, T. Chen, Krüppel-like factor 2 inhibits hepatocarcinogenesis through negative regulation of the Hedgehog pathway, Cancer Sci 110 (2019) 1220–1231. 10.1111/cas.13961.

[46] L.A. Garrett-Sinha, H. Eberspaecher, M.F. Seldin, B. de Crombrugghe, A gene for a novel zinc-finger protein expressed in differentiated epithelial cells and transiently in certain mesenchymal cells, J Biol Chem 271 (1996) 31384–31390. 10.1074/jbc.271.49.31384.

[47] D. Wei, W. Gong, M. Kanai, C. Schlunk, L. Wang, J.C. Yao, T.-T. Wu, S. Huang, K. Xie, Drastic down-regulation of Krüppel-like factor 4 expression is critical in human gastric cancer development and progression, Cancer Res 65 (2005) 2746–2754. 10.1158/0008-5472.CAN-04-3619.

[48] B.D. Rowland, R. Bernards, D.S. Peeper, The KLF4 tumour suppressor is a transcriptional repressor of p53 that acts as a context-dependent oncogene, Nat Cell Biol 7 (2005) 1074–1082. 10.1038/ncb1314.

[49] Z. Xie, Q. Yang, F. Lan, W. Kong, S. Zhao, J. Sun, Y. Yan, Z. Quan, Z. Bai, H. Qing, J. Mao, J. Ni, Cathepsin B deficiency disrupts cortical development via PEG3, leading to depression-like behavior, Commun Biol 8 (2025) 1097. 10.1038/s42003-025-08508-8.

[50] D. Chakroborty, C. Sarkar, R.B. Mitra, S. Banerjee, P.S. Dasgupta, S. Basu, Depleted dopamine in gastric cancer tissues: dopamine treatment retards growth of gastric cancer by inhibiting angiogenesis, Clin Cancer Res 10 (2004) 4349–4356. 10.1158/1078-0432.CCR-04-0059.

[51] C. Sarkar, D. Chakroborty, S. Goswami, H. Fan, X. Mo, S. Basu, VEGF-A controls the expression of its regulator of angiogenic functions, dopamine D2 receptor, on endothelial cells, J Cell Sci 135 (2022) jcs259617. 10.1242/jcs.259617.

[52] D. Chakroborty, S. Goswami, H. Fan, W.L. Frankel, S. Basu, C. Sarkar, Neuropeptide Y, a paracrine factor secreted by cancer cells, is an independent regulator of angiogenesis in colon cancer, Br J Cancer 127 (2022) 1440–1449. 10.1038/s41416-022-01916-1.

[53] S.A. Arnold, L.B. Rivera, J.G. Carbon, J.E. Toombs, C.-L. Chang, A.D. Bradshaw, R.A. Brekken, Losartan slows pancreatic tumor progression and extends survival of SPARC-null mice by abrogating aberrant TGFβ activation, PLoS One 7 (2012) e31384. 10.1371/journal.pone.0031384.

[54] E. Tabatabai, M. Khazaei, F. Asgharzadeh, S.E. Nazari, N. Shakour, H. Fiuji, A. Ziaeemehr, A. Mostafapour, M.R. Parizadeh, M. Nouri, S.M. Hassanian, F. Hadizadeh, G.A. Ferns, M. Rahmati, F. Rahmani, A. Avan, Inhibition of angiotensin II type 1 receptor by candesartan reduces tumor growth and ameliorates fibrosis in colorectal cancer, EXCLI J 20 (2021) 863–878. 10.17179/excli2021-3421.

[55] N. Fedchenko, J. Reifenrath, Different approaches for interpretation and reporting of immunohistochemistry analysis results in the bone tissue - a review, Diagn Pathol 9 (2014) 221. 10.1186/s13000-014-0221-9.

[56] P. Agyakwa, J. Dai, J. Li, B. Mouawad, L. Yang, M. Corfield, C.M. Johnson, Three-dimensional damage morphologies of thermomechanically deformed sintered nanosilver die attachments for power electronics modules, J Microsc 277 (2020) 140–153. 10.1111/jmi.12803.

[57] M.F. Siqueira, S. Flowers, R. Bhattacharya, D. Faibish, Y. Behl, D.N. Kotton, L. Gerstenfeld, E. Moran, D.T. Graves, FOXO1 modulates osteoblast differentiation, Bone 48 (2011) 1043–1051. 10.1016/j.bone.2011.01.019.

[58] M. Junking, T. Rattanaburee, A. Panya, I. Budunova, G. Haegeman, P.-T. Yenchitsomanus, Anti-Proliferative Effects of Compound A and Its Effect in Combination with Cisplatin in Cholangiocarcinoma Cells, Asian Pac J Cancer Prev 21 (2020) 2673–2681. 10.31557/APJCP.2020.21.9.2673.

[59] C.-C. Liang, A.Y. Park, J.-L. Guan, In vitro scratch assay: a convenient and inexpensive method for analysis of cell migration in vitro, Nat Protoc 2 (2007) 329–333. 10.1038/nprot.2007.30.

[60] A. Albini, Y. Iwamoto, H.K. Kleinman, G.R. Martin, S.A. Aaronson, J.M. Kozlowski, R.N. McEwan, A rapid in vitro assay for quantitating the invasive potential of tumor cells, Cancer Res 47 (1987) 3239–3245.

[61] J.D. Nelson, O. Denisenko, K. Bomsztyk, Protocol for the fast chromatin immunoprecipitation (ChIP) method, Nat Protoc 1 (2006) 179–185. 10.1038/nprot.2006.27.

[62] Z. Wang, M. Gerstein, M. Snyder, RNA-Seq: a revolutionary tool for transcriptomics, Nat Rev Genet 10 (2009) 57–63. 10.1038/nrg2484.

[63] D. Parkhomchuk, T. Borodina, V. Amstislavskiy, M. Banaru, L. Hallen, S. Krobitsch, H. Lehrach, A. Soldatov, Transcriptome analysis by strand-specific sequencing of complementary DNA, Nucleic Acids Res 37 (2009) e123. 10.1093/nar/gkp596.

64. [64] S. Anders, W. Huber, Differential expression analysis for sequence count data, Genome Biol 11 (2010) R106. 10.1186/gb-2010-11-10-r106.

[65] A. Subramanian, P. Tamayo, V.K. Mootha, S. Mukherjee, B.L. Ebert, M.A. Gillette, A. Paulovich, S.L. Pomeroy, T.R. Golub, E.S. Lander, J.P. Mesirov, Gene set enrichment analysis: a knowledge-based approach for interpreting genome-wide expression profiles, Proc Natl Acad Sci U S A 102 (2005) 15545–15550. 10.1073/pnas.0506580102.

[66] D. Tang, M. Chen, X. Huang, G. Zhang, L. Zeng, G. Zhang, S. Wu, Y. Wang, SRplot: A free online platform for data visualization and graphing, PLoS One 18 (2023) e0294236. 10.1371/journal.pone.0294236.

[67] D.S. Chandrashekar, B. Bashel, S.A.H. Balasubramanya, C.J. Creighton, I. Ponce-Rodriguez, B.V.S.K. Chakravarthi, S. Varambally, UALCAN: A Portal for Facilitating Tumor Subgroup Gene Expression and Survival Analyses, Neoplasia 19 (2017) 649–658. 10.1016/j.neo.2017.05.002.

[68] D.S. Chandrashekar, S.K. Karthikeyan, P.K. Korla, H. Patel, A.R. Shovon, M. Athar, G.J. Netto, Z.S. Qin, S. Kumar, U. Manne, C.J. Creighton, S. Varambally, UALCAN: An update to the integrated cancer data analysis platform, Neoplasia 25 (2022) 18–27. 10.1016/j.neo.2022.01.001.

[69] Z. Tang, C. Li, B. Kang, G. Gao, C. Li, Z. Zhang, GEPIA: a web server for cancer and normal gene expression profiling and interactive analyses, Nucleic Acids Res 45 (2017) W98–W102. 10.1093/nar/gkx247.

