## Supplementary file for "Angiotensin II Type 1 Receptor Blockade Inhibits Gastric Cancer Metastasis Through Tight Junction Restoration"

[Sooraj Kakkat](https://pubmed.ncbi.nlm.nih.gov/?sort=date&size=200&term=Kakkat+S&cauthor_id=37108147) ^1,2^, Prabhat Suman ^1,2^, Sandeep Goswami ^1,2^, Brusi Kola ^1,2^, Kira A Bruno ^2^, Wendy L Frankel ^3,4^, Sujit Basu ^3,4,5^, Elba A Turbat-Herrera^1,2^, Veronica Ramirez-Alcantara^2^, Martin J Heslin^2^, Joel F Andrews^2^, Paramahansa Pramanik ^6^, Chandrani Sarkar ^1,2, 7^, Debanjan Chakroborty ^1,2, 7*^

^1^ Department of Pathology, University of South Alabama, Mobile, AL 36617, USA

^2^ Mitchell Cancer Institute, University of South Alabama, Mobile, AL 36604, USA

^3^ Department of Pathology, The Ohio State University, Columbus, OH, USA

^4^ Comprehensive Cancer Center, The Ohio State University, Columbus, OH, USA

^5^ Department of Medical Oncology, The Ohio State University, Columbus, OH, USA
^6^ Department of Mathematics and Statistics, University of South Alabama, Mobile, AL	36688, USA

^7^ Department of Biochemistry and Molecular Biology, University of South Alabama, Mobile, AL 36688, USA

^*^Correspondence:

Debanjan Chakroborty, PhD

Assistant Professor of Pathology

University of South Alabama

+1-251-445-8403

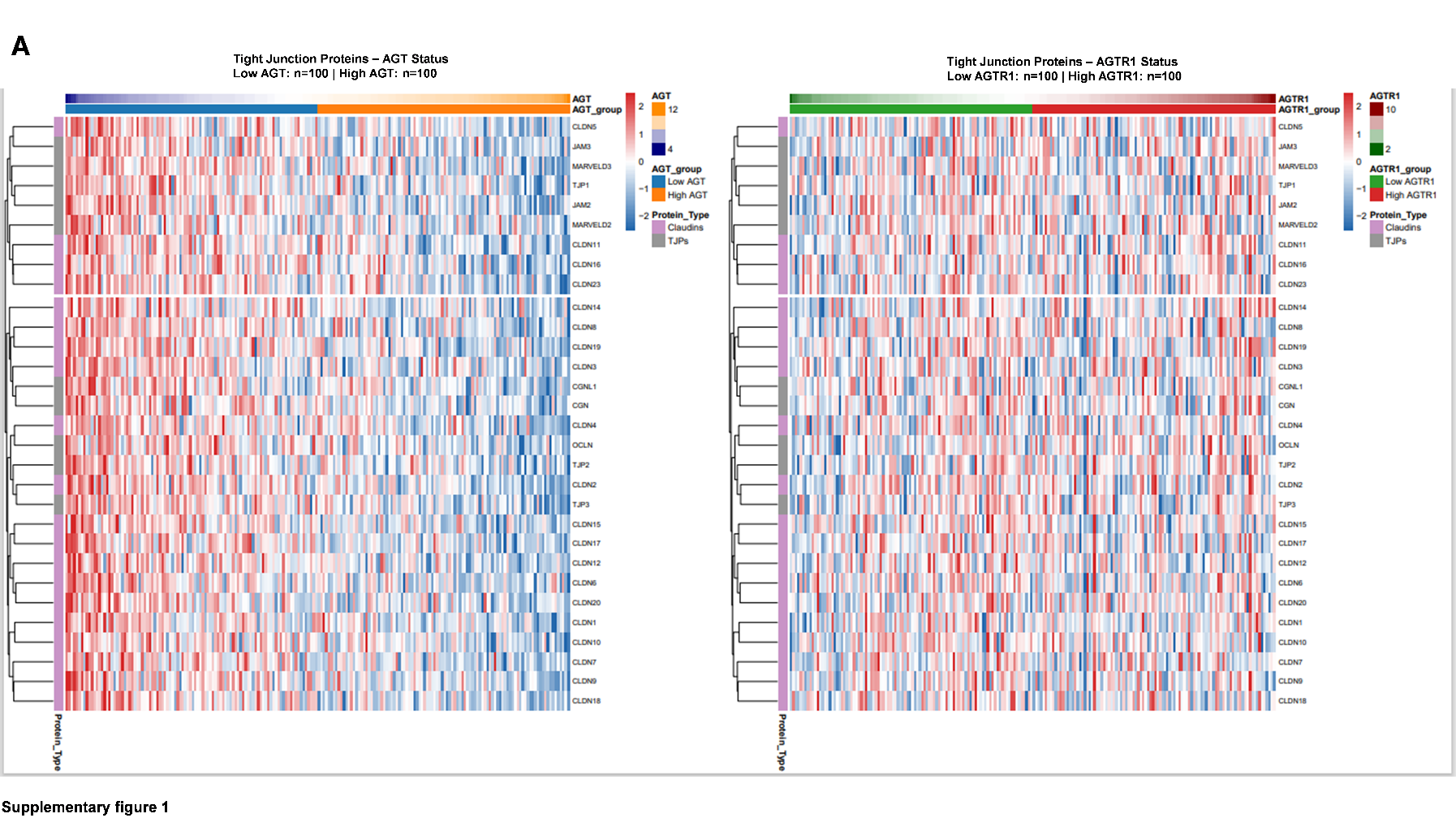

**Fig. S1. Tight junction gene expressions and their association with *AGT* and *AGTR1* in TCGA-STAD. (A)** The heatmaps display Z-score-scaled expression of TJ-related genes across TCGA-STAD tumors (n = 100 per group for each gene). The column represents individual tumors, and rows represent the TJ-related genes. The Z score blue, white, and red denotes the relative under- to over-expression across the cohort. The top bars indicate the continuous *AGT* or *AGTR1* expression and their status (Low vs High). Low *AGT* tumors generally have higher TJ gene expression, while high *AGT* tumors show downregulation of these TJ genes. AGT, angiotensin II; AGTR1, angiotensin II receptor type 1.

| 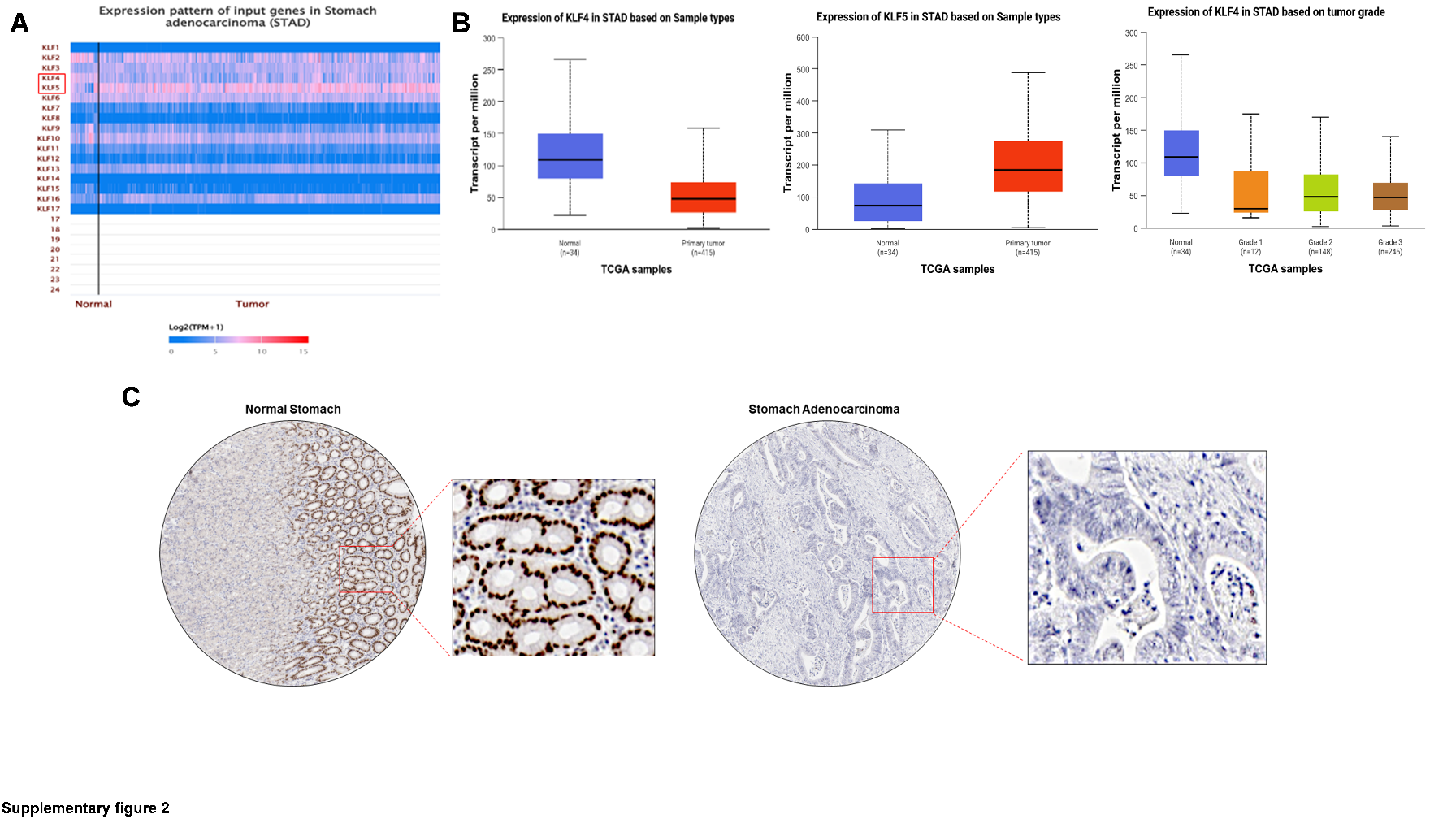  **Fig. S2. KLF4 downregulation in GC is associated with poor patient prognosis.** (**A)** Heatmap generated using TCGA transcriptomic data shows the expression of Kruppel-like factor (*KLF*) family genes in normal gastric (*n* = 34) and primary stomach adenocarcinoma tissues (STAD) (*n* = 415). Data indicate a consistent change in *KLF4* (downregulation or loss) and *KLF5* (upregulation) expression in human GC tissues among 17 *KLFs.* (**B)** Box plots show significantly decreased *KLF4* and increased *KLF5* transcript levels in STAD tumors compared to normal tissues (data from TCGA). It also shows a stage-wise decrease in KLF4 expression in STAD. (**C)** Representative IHC images from The Human Protein Atlas reveal a significant loss of KLF4 in malignant gastric cancer tissue. GC, gastric cancer; STAD, stomach adenocarcinoma; TCGA, The Cancer Genome Atlas; IHC, immunohistochemistry. | |
| --- | --- |
|  | 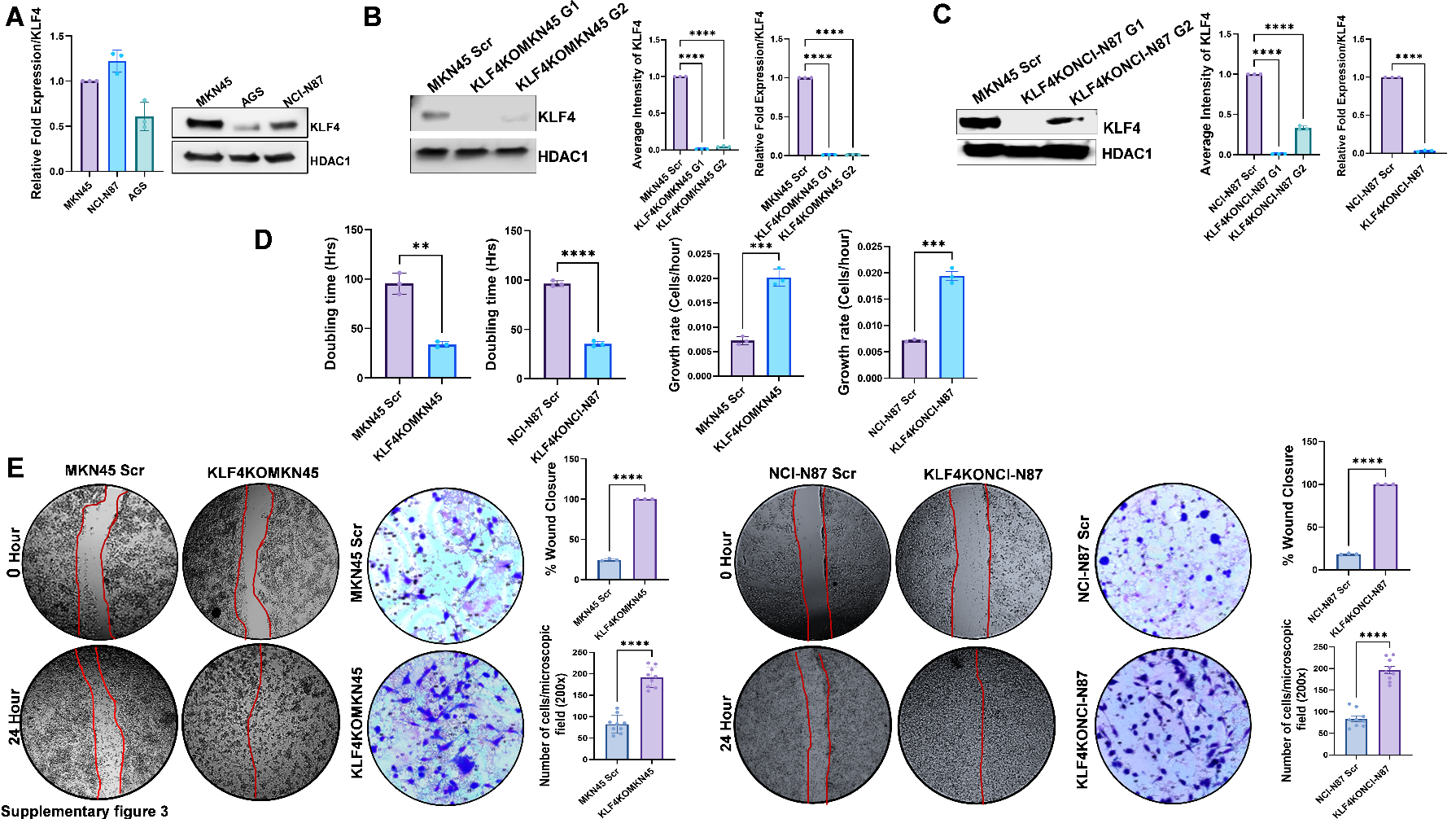  **Fig. S3.** **Loss of KLF4 promotes cell proliferation and metastasis of GC cells. (A)** Representative qRT-PCR and western blot images show KLF4 expression in different GC cell lines.  Data represented as mean±SD. (**B and C)** Western blot and qRT-PCR analysis confirm the deletion of KLF4 in MKN45 and NCI-N87 cells after CRISPR/Cas9-mediated knockout. For mRNA quantification, KLF4 was normalized to the housekeeping gene GAPDH. Densitometric quantification of KLF4 protein levels normalized to GAPDH from the western blots.  Significance was determined by 1-way ANOVA with Dunnett’s multiple comparisons test and 2-tailed *t* test (*n* = 3) (**D).** A significant decrease in cell doubling time and an increase in cell proliferation are observed in KLF4KOMKN45 and KLF4KONCI-N87 cells compared to their scrambled control cells (Scr). Significance was determined by a 2-tailed *t* test *(n = 3).* (**E)** KLF4 knockout MKN45 and NCI-N87 cells (KLF4KOMKN45 and KLF4KONCI-N87) show markedly enhanced cell migration *in vitro*, as demonstrated by wound healing assays at 0 h and 24 h. Magnification, 40X. Transwell invasion assays revealed significantly increased invasive capacity in *KLF4*-deleted cells compared to scrambled cells. Magnification, 200X. Quantification of migrated and invaded cells is shown in bar graphs (right panels). Data represented as mean±SEM*.* Significance was determined by a 2-tailed t-test (migration, *n* = 3; invasion, *n* =9). **P <* 0.05*; **P* < 0.01*; ***P* < 0.001*;****P* < 0.0001. GC, gastric cancer; Scr, scrambled control. |

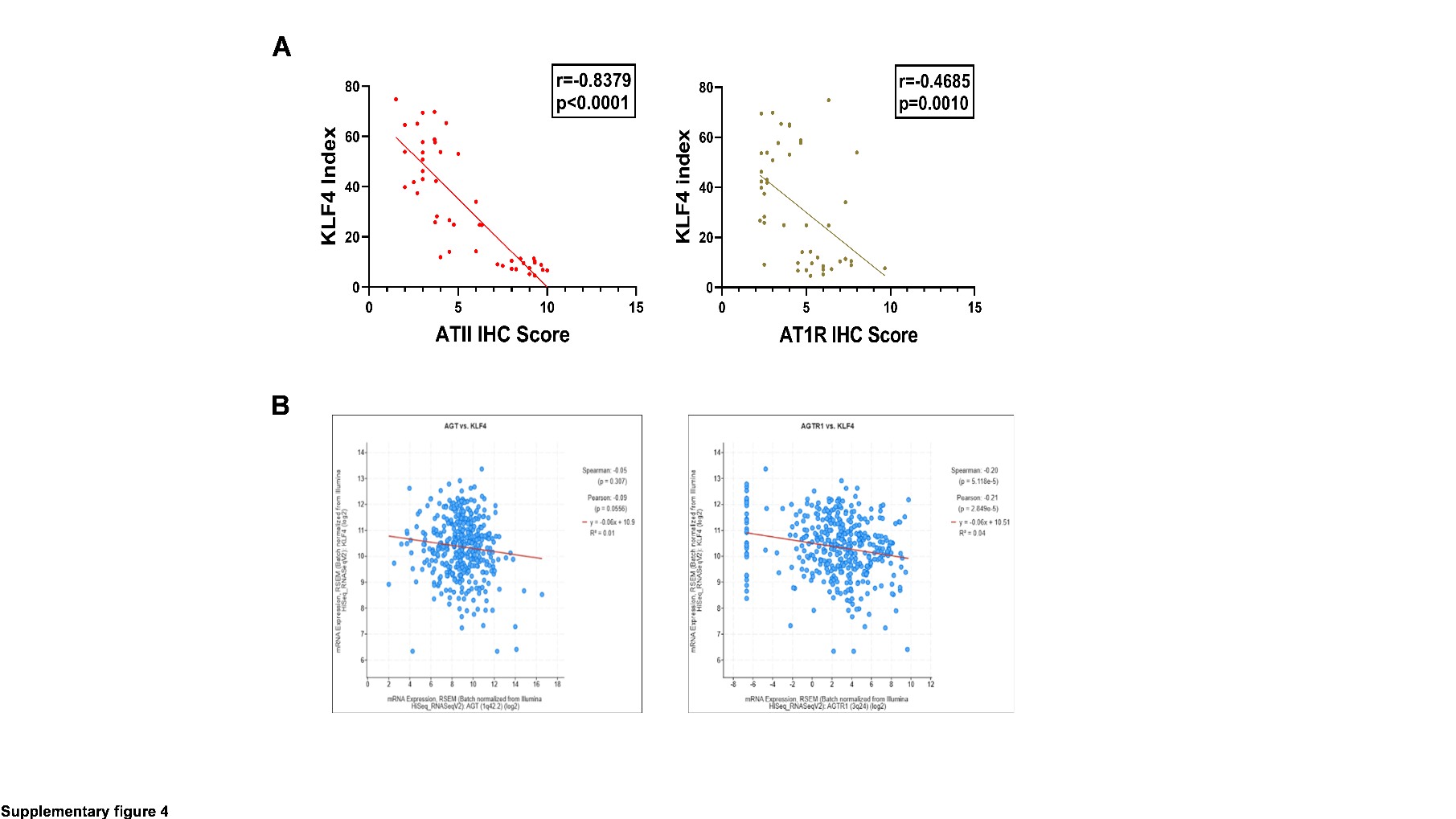

**Fig. S4. ATII/AT1R expression is negatively associated with expression of KLF4 in GC. (A)** The quantitative correlation analysis shows a significant negative correlation between the average IHC score for ATII/AT1R expression and the number of KLF4-positive cells. Correlation was assessed using Pearson’s correlation test; results are displayed in the graph (*n =* 46). **(B)** cBioPortal data show that elevated mRNA expression of *AGT* and *AGTR1* is negatively correlated with *KLF4* expression in GC. **P* < 0.05*; **P* < 0.01*; ***P <* 0.001*; ****P* < 0.0001. GC, gastric cancer; TJ, tight junction, AGT, angiotensin II; AGTR1, angiotensin II receptor type 1.

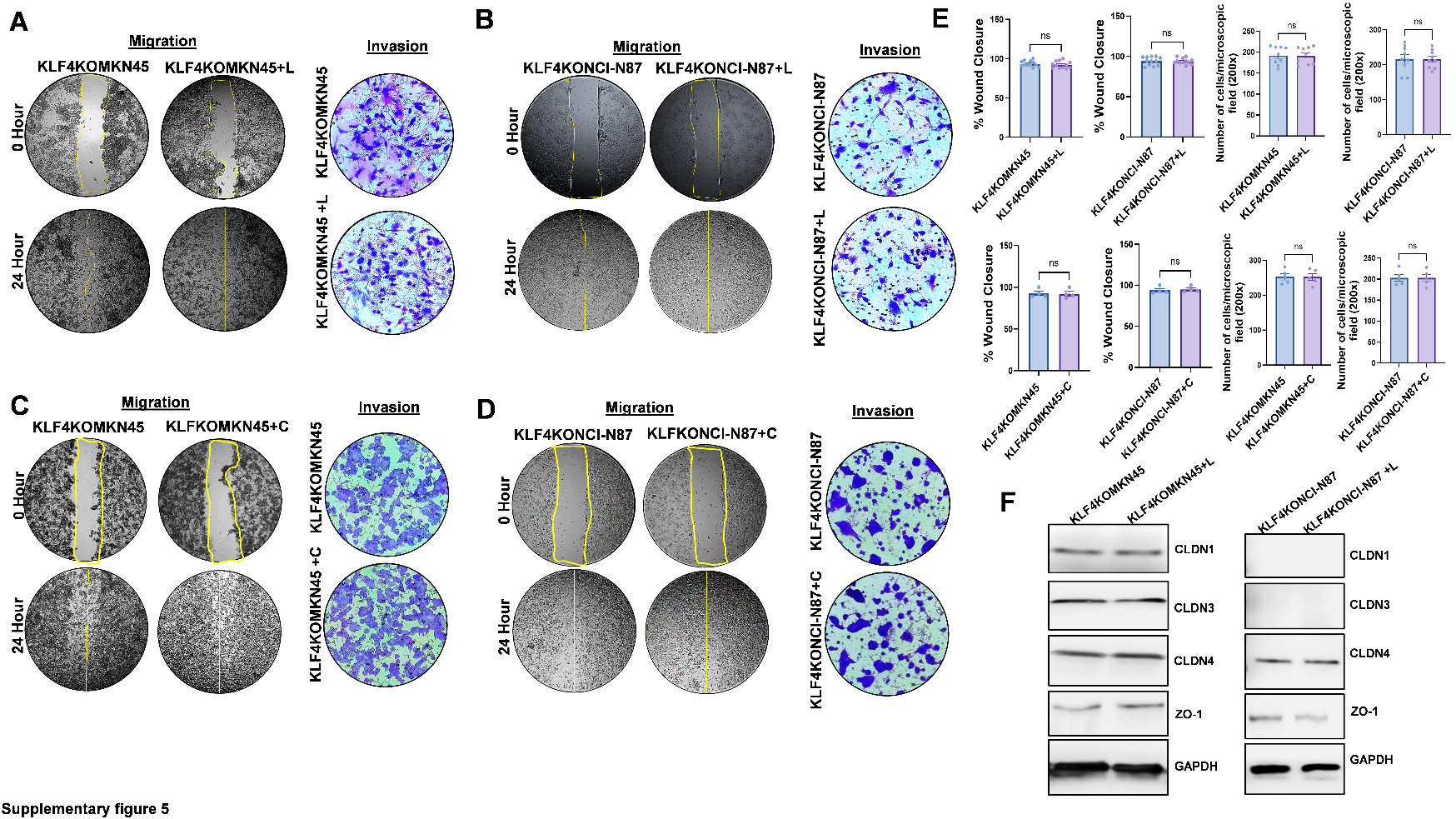

**Fig**. **S5.** **AT1R inhibition fails to restore the expression of TJ proteins or inhibit the metastatic capacity of KLF4-knockout GC cells.** **(A-D)** Wound closure and trans well assays show that neither losartan nor candesartan treatment significantly alters the migratory or invasive potential of KLF4KOMKN45 and KLF4KONCI-N87 cells compared to untreated cells (migration, Magnification 40X; invasion, Magnification 200X). **(E)** The percentage of wound closure and cell-invasive index after losartan or candesartan treatment is shown as bar diagrams (data as mean±SEM), with significance assessed by 2-tailed t-test (*n* = 12 for losartan wound closure, *n* = 9 for losartan invasion, *n* = 4 for candesartan wound closure, *n* = 5 for candesartan invasion). **(F)** Western blot analysis shows no significant change in levels of TJ proteins Claudin 1, 3, 4, and Zonula occludens 1 in losartan-treated cells. *ns*: no significance. GC, gastric cancer; TJs, tight junctions.

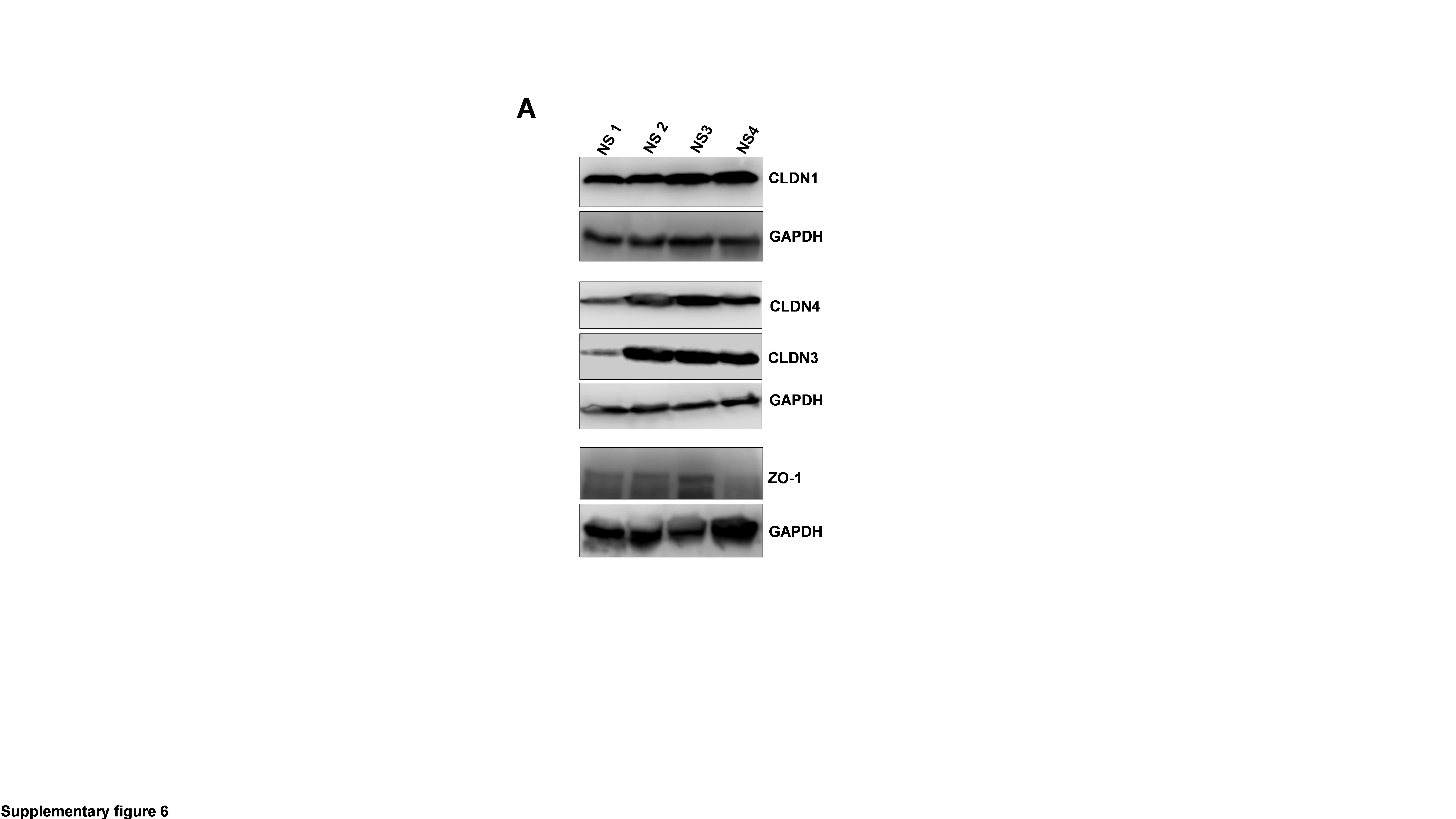

**Fig. S6. Western blot analysis of TJ protein expression in normal mouse stomach tissue.** **(A)** Representative western blot data show the expression levels of Claudin 1, 3, 4, and Zonula occludens 1 in normal mouse stomach tissue. GAPDH was used as a loading control. TJ, tight junction ; NS, normal stomach.

**Table S1.** List of antibodies used for immunohistochemistry, western-blot and immunofluorescence analyses.

| **Primary Antibody** | **Dilution** | | | **Source** | **Catalogue No** |
| --- | --- | --- | --- | --- | --- |
|  | **Western Blot** | **Immunohistochemistry** | **Immunofluorescence** |  |  |
| Angiotensin II/III Antibody (Ang II E7 (BGN/KA/4L)) | 1:800 | 1:100 | - | Novus Biologicals, CO | NB 100-62346 |
| Anti-Angiotensin II Type 1 Receptor antibody | 1:1000 | 1:50 | - | Abcam, MA | ab124505 |
| Anti-KLF4 antibody | 1:500 | 1:100 | 1:800 | Abcam, MA | ab216968 |
| KLF4 Antibody | 1:1000 | 1:100 | 1:800 | Cell Signaling Technology, Danvers, MA | 4038S |
| CLDN1 monoclonal antibody (M01), clone 1C5-D9 | 1:1000 | 1:100 | 1:600 | Abnova, Taipei, Taiwan | H00009076-M01 |
| Claudin-3 Antibody (OTI1E7) | 1:1000 | 1:100 | - | Novus Biologicals, CO | NBP2-46299 |
| Anti-Claudin 4 antibody | 1:1000 | 1:100 | 1:600 | Abcam, MA | ab53156 |
| Tight Junction Protein 1 Antibody | 1:1000 | 1:100 | 1:500 | Novus Biologicals, CO | NBP2-80141 |

**Table S2.** Sequences of primers used in qRT-PCR

| **Primer Name** | **Accession &**  **Definition** | **Primer Pair Sequence 5’-**  **3’(Sense/Antisense)** | **Fragment**  **Size (bp)** |
| --- | --- | --- | --- |
| *KLF4* | NM_001314052.2 | CATCTCAAGGCACACCTGCGAA  TCGGTCGCATTTTTGGCACTGG | 156 |
| *CLDN1* | NM_021101.5 | GTCTTTGACTCCTTGCTGAATCTG  CACCTCATCGTCTTCCAAGCAC | 144 |
| *CLDN3* | NM_001306.4 | GCCTTCATCGGCAGCAACATCA  AGCGAGTCGTTACACCTTGCACT | 111 |
| *CLDN4* | NM_001305.5 | AGTGCAAGGTGTACGACTCGCT  CGCTTTCATCCTCCAGGCAGTT | 150 |
| *TJP1* | XM_054330054.1 | GTCCAGAATCTCGGAAAAGTGCC  CTTTCAGCGCACCATACCAACC | 132 |
| *AGT* | NM_001384479.1 | TGGACAGCACCCTGGCTTTCAA  ACACTGAGGTGCTGTTGTCCAC | 111 |
| *AGTR1* | NM_000685.5 | CAGCGTCAGTTTCAACCTGTACG  GCAGGTGACTTTGGCTACAAGC | 130 |
| *GAPDH* | NM_001289746.2 | GTCTCCTCTGACTTCAACAGCG  ACCACCCTGTTGCTGTAGCCAA | 131 |

**Table S3.** Sequences of primers used in ChIP assay

| **Primer Name** | **Primer Pair Sequence 5’-**  **3’(Sense/Antisense)** | **Location (nt)** | **Fragment**  **Size(bp)** |
| --- | --- | --- | --- |
| *CLDN1* | GCGAGAAGATCCACGAGAGA  GATTTAAAGCAGCTCCGCCC | chr3:190322467-190322657 | 191 |
| *CLDN3* | TGGGCCCAGCAATTCCTG  TTGACACGGCTTCTCTCTCC | chr7:73770586-73770819 | 234 |
| *CLDN4* | ATCAAGGTGGGGTTTCTCGT  GTTCACAGGTTAAGGCCAGG | chr7:73830553+73830733 | 181 |
| *TJP1* | CTAGGTGGCCTTCCTGCATTT  TTTGCCCTTCCCCTATACTAACG | chr15:27489152+27489394 | 243 |
